# Co-option and Detoxification of a Phage Lysin for Housekeeping Function

**DOI:** 10.1101/418723

**Authors:** Amelia M. Randich, David T. Kysela, Cécile Morlot, Yves V. Brun

**Affiliations:** Indiana University, Bloomington, IN USA; Institut de Biologie Structurale (IBS), Université Grenoble Alpes, CEA, CNRS, France

**Keywords:** Lysozyme, GH24, Prophage domestication, Bacterial evolution, Alphaproteobacteria, *Caulobacter*, *Asticcacaulis*

## Abstract

Temperate phages constitute a potentially beneficial genetic reservoir for bacterial innovation despite being selfish entities encoding an infection cycle inherently at odds with bacterial fitness. These phages integrate their genomes into the bacterial host during infection, donating new, but deleterious, genetic material: the phage genome encodes toxic genes, such as lysins, that kill the bacterium during the phage infection cycle. Remarkably, some bacteria have exploited the destructive properties of phage genes for their own benefit by co-opting them as toxins for functions related to bacterial warfare, virulence, and secretion. However, do toxic phage genes ever become raw material for functional innovation? Here we report on a toxic phage gene whose product has lost its toxicity and has become a domain of a core cellular factor, SpmX, throughout the bacterial order Caulobacterales. Using a combination of phylogenetics, bioinformatics, structural biology, cell biology, and biochemistry, we have investigated the origin and function of SpmX and determined that its occurrence is the result of the detoxification of a phage peptidoglycan hydrolase gene. We show that the retained, attenuated activity of the phage-derived domain plays an important role in proper cell morphology and developmental regulation in representatives of this large bacterial clade. To our knowledge, this is the first observation of phage gene domestication in which a toxic phage gene has been co-opted for a housekeeping function.

## Introduction

Understanding how new genes arise is key to studying the forces that drive diversity and evolution. Although horizontal gene transfer (HGT) is widely regarded as an important mechanism for the exchange of existing genes among bacteria, mobile genetic elements can transfer exogenous genetic material that gives rise to novel genes. These new genes can provide the basis for evolving new traits and propelling evolutionary transitions [1,2]. Temperate bacteriophages mediate genetic transfer by integrating their genomes into bacterial hosts [3–6]. These integrated tracts of genes, called prophages, remain dormant until induced by various signals to produce phage particles and phage-encoded proteins that lyse the cell. In many cases, prophages contain genes that benefit the host, promoting retention of prophages in many bacterial lineages, even after mutations have inactivated the prophages and they no longer are capable of producing phage [7–9]. Accumulation of host-specific beneficial mutations in prophages has been referred to as “domestication.” Many so-called domesticated segments of inactivated prophages unexpectedly contain lytic and phage structural genes, which would intuitively be of little use or even detrimental to the bacterial host [7]. These genes are often used by the bacteria as weapons against competing bacteria and eukaryotic hosts [10–14]. In contrast, we have identified an instance in which a toxic phage gene has not been repurposed as a weapon, but has rather been incorporated into a bacterial housekeeping factor, *spmX*. Here, we report that SpmX resulted from an ancient domestication event at the root of the alphaproteobacterial order Caulobacterales, in which co-option and detoxification of a toxic phage gene gave rise to a novel bacterial gene with roles in developmental regulation and morphogenesis.

SpmX was first identified as a developmental regulator in the model organism *Caulobacter crescentus* [15]. Like most members of Caulobacterales, stalked *C. crescentus* cells divide asymmetrically to produce a “mother” cell with a polar stalk and a motile “daughter” or “swarmer” cell with a polar flagellum. The *Caulobacter* developmental cycle depends on strict coordination of cell growth, chromosome replication and segregation, and division by various regulatory proteins that differ in localization and timing [16]. This network depends on regulatory phospho-signaling factors localized and regulated by polar scaffolds. SpmX is one scaffold that localizes at the old pole during the swarmer-to-stalked cell transition and recruits and potentially activates the histidine kinase DivJ [15]. Intriguingly, SpmX is required for stalk synthesis initiation and elongation in the closely related *Asticcacaulis* species *A. excentricus* and *A. biprosthecum* [17]. Therefore, this gene appears to have evolved multiple roles within this family of dimorphic, stalked bacteria.

Perplexingly, SpmX contains an N-terminal putative phage muramidase domain that is generally toxic to bacteria. This enzymatic domain is typically used by phage to lyse bacteria and release infectious phage particles during the lytic cycle. As a part of SpmX, this domain is critical for SpmX’s role in both developmental regulation and stalk biogenesis. In particular, the muramidase domain is necessary for proper localization of SpmX in both *C. crescentus* [15,18] and the *Asticcacaulis* genus [17]. Various studies have shown that SpmX localizes with the polar scaffold PopZ in *C. crescentus* [19,20], and this interaction occurs entirely through the muramidase domain [18]. A previous study was unable to measure enzymatic activity from purified *C. crescentus* SpmX muramidase domain, and consequently concluded that the domain was repurposed for protein interactions and oligomeric assembly [18]. However, given the remarkable sequence similarity of the SpmX muramidase domain to functional phage lysozymes, including the canonical catalytic glutamate, it seems unlikely that its enzymatic activity is entirely lost. Why would this domain be so highly conserved if its new function were merely for non-essential protein-protein interactions?

To better characterize the SpmX muramidase domain and the constraints underlying its conservation, we performed an in-depth bioinformatics study of more than 60 available SpmX genes together with structural determination, biochemical analysis, and comparative cell biology between *Caulobacter* and *Asticcacaulis.* We show that SpmX arose prior to the diversification of Caulobacterales, a large class of stalked bacteria. We establish that the SpmX muramidase domain is a close relative of GH24 (T4-like) autolysin/endolysins that have been laterally exchanged throughout Proteobacteria via prophages. We find that the SpmX domain is still an active muramidase, albeit with attenuated ancestral phage activity, consistent with its remodeled active cleft. Finally, we demonstrate that the enzymatic activity is necessary for SpmX function in three representative species. We conclude that, close to the time of the genesis of the full-length *spmX* gene, this co-opted domain accumulated mutations that attenuated its hydrolytic activity on peptidoglycan and detoxified it for bacterial use. To our knowledge, this is the first case of phage gene domestication in which a toxic phage gene has been incorporated into a new bacterial gene with housekeeping function.

## Results

### The SpmX muramidase domain was co-opted from prophage in an early Caulobacterales ancestor

Our first step in characterizing the muramidase domain of SpmX was to determine the prevalence of SpmX and its homologues in the bacterial domain. Simple pBLAST analysis revealed that SpmX, as defined by its three-part architecture with an N-terminal muramidase domain, a charged and proline-rich intermediate domain, and two C-terminal transmembrane (TM) segments (**Fig. 1A**), is taxonomically constrained to Caulobacterales and one member of its sister taxa, Parvularculales. It is conserved as a single-copy gene in all sequenced members (**Table S1**). In all 69 identified *spmX* orthologues, the muramidase domains exhibit high amino acid sequence conservation (**Fig. S1**), the TM segments moderate sequence conservation, and the intermediate domains high variability in length and composition among genera (**Fig. 1A**). Apart from these SpmX orthologues, BLAST searches using SpmX only returned hits for the muramidase domain. These hits came from Gram-negative bacterial genomes that span the entire bacterial domain, particularly proteobacteria, and from viral genomes. Most of these bacterial genes are likely to be in prophage regions, as evidenced by their position in long runs of predicted prophage genes. We did not detect sequences homologous to SpmX TMs in our search, although we occasionally detected homologous phage muramidase domains fused to other, non-homologous TM segments.

**Figure 1.**
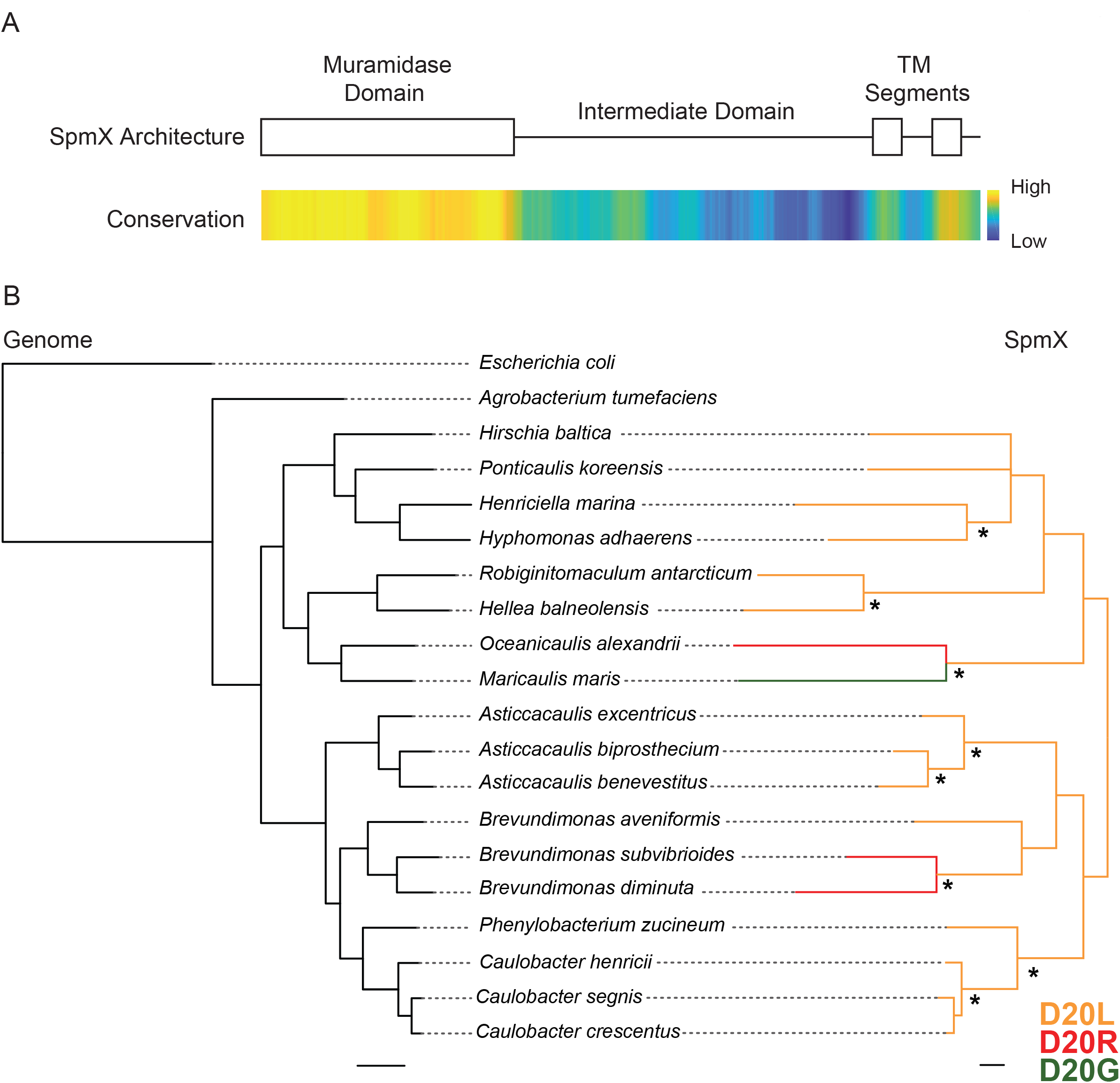
The SpmX is vertically inherited in Caulobacterales. **(A)** Schematic of SpmX architecture, including the conserved muramidase domain (see **Fig. S1** for alignments), the variable intermediate domain, and two C-terminal transmembrane (TM) segments. Bar indicates amino acid sequence conservation among *spmX* alleles. **(B)** Phylogenetic trees of representative species from Caulobacterales and other Alphaproteobacteria for concatenated housekeeping gene alignments (left) and for SpmX (right), with branch colors indicating the amino acid identity at position 20 of SpmX (D20L in yellow, D20R in red, and D20G in green). The concatenated housekeeping tree is fully supported with posterior probability of 1.0 for all clades. Asterisks indicate clades in the SpmX tree with posterior probabilities > 0.95.

Consistent with finding close relatives of SpmX muramidase in prophages, NCBI’s Conserved Domain Database (CDD) tool [21,22] clustered SpmX muramidase with glycoside hydroloase (GH) 24 lysozymes in the autolysin/endolysin class (**Fig. S2)**. These enzymes are closely related to classical phage T4 lysozyme-like (T4L-like) peptidoglycan hydrolases. Phages generally use lysozymes of this class to cleave peptidoglycan and lyse cells during the lytic cycle. These lysozymes are distinct from the other well-known phage Lambda lysozyme-like lytic transglycosylases, which are related to known housekeeping bacterial hydrolases with roles in cell growth and division. Lytic transglycosylases are also assigned to the GH24 group but share no sequence similarity with and catalyze a different peptidoglycan cleaving reaction than T4L-like muramidases [23–25]. Thus although core bacterial genomes encode peptidoglycan hydrolases, the muramidase domain of SpmX is most closely related to peptidoglycan hydrolases encoded by prophages.

Unlike its close relatives that have been transferred horizontally through the bacterial domain via prophage, the SpmX muramidase domain coding region has been inherited vertically as part of the *spmX* gene in Caulobacterales. The SpmX gene tree mirrors the phylogeny of Caulobacterales from concatenated gene alignments (**Fig. 1B**). None of the SpmX genes appear in tracks of prophage genes, and the genomic context of *spmX* appears to be well maintained in members of Caulobacterales, with the gene occurring between a putative Mg^2+^ transporter and a putative isovaleryl-CoA dehydrogenase in nearly all species. Together, these findings suggest that SpmX muramidase is a peptidoglycan hydrolase derived from prophage that has become a domain in a bacterial protein that is no longer within prophage and is under direct cellular control. It likely fused with the intermediate and TM domains in a common ancestor of Parvularculales and Caulobacterales. The vertical transmission of SpmX and strong sequence conservation of the muramidase domain suggests an important function in members of Caulobacterales.

### The SpmX muramidase domain retains the canonical GH24 motif but contains mutations in the catalytic cleft known to inactivate phage lysozymes

To determine if critical enzymatic residues in SpmX muramidase were conserved, we compared the amino acid sequences of SpmX and other lysozymes. By definition, lysozymes catalyze the hydrolysis of β1,4-linked glycosidic bonds in peptidoglycan and chitin [25]. This superfamily includes at least seven distinct groups that are unrelated by sequence similarity but share a common fold in which the catalytic Glu and the beta-hairpin motif in the N-terminal lobe form the catalytic cleft by packing against the C-terminal lobe (**Fig. 2A)** [26]. This beta-hairpin, or GH motif, contains family-specific residues critical for enzyme activity in all members of the lysozyme superfamily [26].

**Figure 2.**
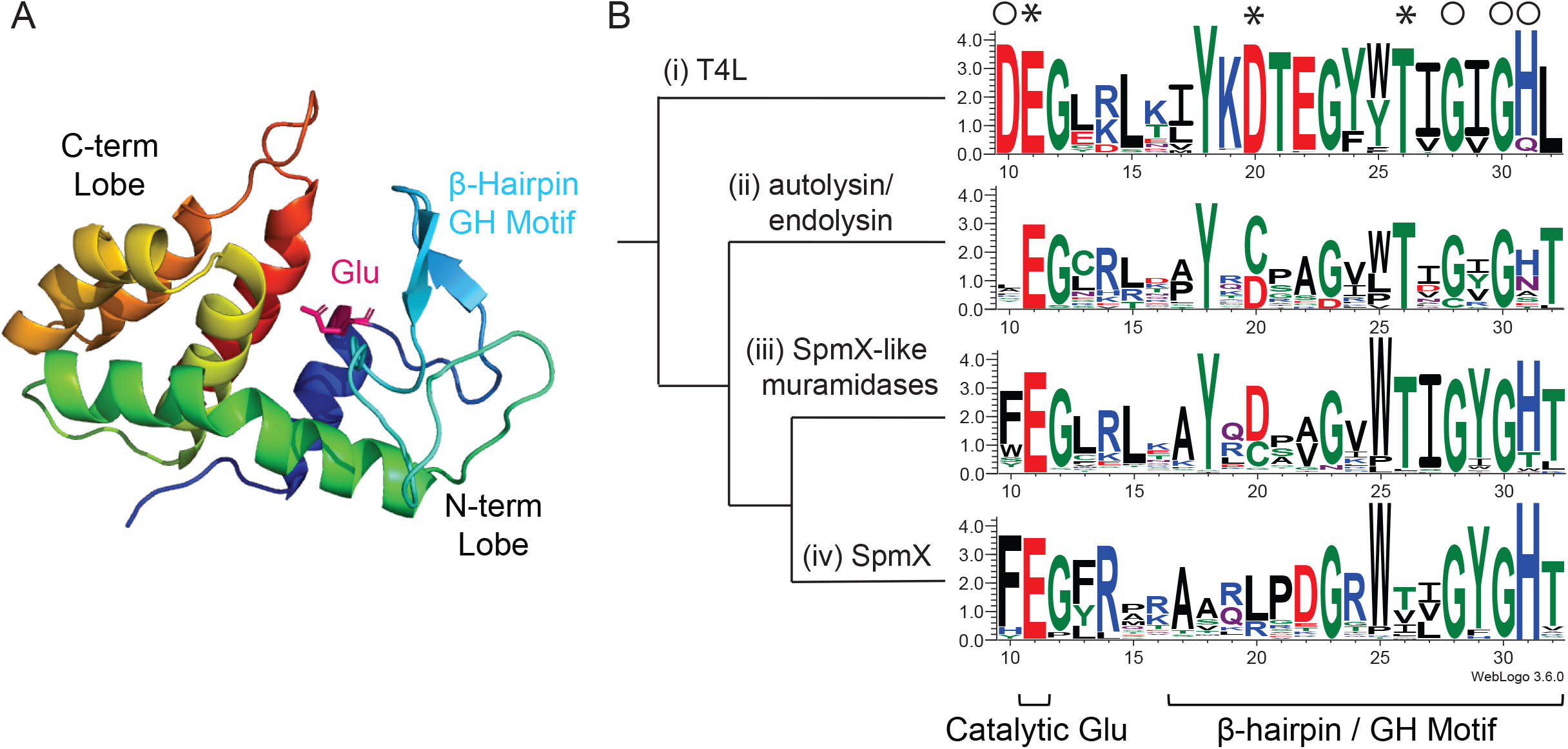
The SpmX muramidase domain retains the canonical GH motif but contains inactivating mutations in the catalytic cleft. **(A)** P22 lysozyme (PDB 2ANX) as a model lysozyme colored with rainbow gradient from blue N-terminus to red C-terminus. The catalytic glutamate appears in fuchsia and the GH beta-hairpin in light blue. **(B)** HMM logos of GH lysozymes made using WebLogo 3 [53]. Logos were constructed from protein sequences of **(i)** T4 lysozyme-like genes (n = 94), **(ii)** representative autolysins/endolysins from the Conserved Domain Database including P22 lysozyme (n = 20) but excluding SpmX genes, **(iii)** closest BLAST hits from non-SpmX muramidases (n = 60), and **(iv)** SpmX muramidases (n = 66), and organized in a cladogram to resemble the sequence cluster tree diagram in **Figure S2**. Amino acids are color-coded according to chemical properties, with uncharged polar residues in green, neutral residues in purple, basic residues in blue, acidic residues in red, and hydrophobic residues in black. The height of each letter is proportional to the relative frequency of a given identity and the height of the stack indicates the sequence conservation at that position. T4L numbering is used for ease of comparison. Asterisks mark positions critical for enzymatic activity and open circles mark positions associated with GH motif stability [26,27].

We compared the GH motif of SpmX to lysozymes from the T4L and endolysin/autolysin classes with varying degrees of relatedness. **Figure 2B** shows the amino acid conservation in the GH motif of T4L-like, autolysin/endolysin, closely related non-SpmX muramidase, and SpmX muramidase protein sequences. Because the autolysin/endolysin class and the closest non-SpmX relatives are likely to be active phage enzymes, highly conserved residues shared by these groups with T4L delineate positions that are evolutionarily constrained for phage lysozyme activity and stability. For example, D10 is not conserved outside of T4L-like enzymes because the autolysin/endolysin class does not have a salt bridge between D10 and the C-terminal lobe [27]. On the other hand, all of the putative phage sequences (**Fig. 2B(i-iii)**) conserve the “catalytic triad” of T4 lysozymes: the catalytic residue E11 and active site residues D20 and T26. While the exact roles of D20 and T26 are not clear, they are posited as being critical for effective catalysis [26–29]. Position D20 is very sensitive to mutation, with only substitutions D20C/A retaining the hydrolytic activity of T4L or P22 phage lysozymes [30]; these substitutions are tellingly well represented amongst the putative phage sequences. Remarkably, SpmX muramidase domains demonstrate strong conservation of residues required for T4L stability and creating the GH fold, but low conservation of residues associated with catalysis, with the exception of the main catalytic residue, E11 (**Fig. 2B(iv)**). The majority of SpmX genes contain the mutation D20L/R, both of which reduced T4L activity to less than 3% of WT in previous studies [27] and which are distinctly unrepresented in the other phage muramidases. Moreover, the T26 position no longer appears to be under constraint in SpmX. The conservation of the GH motif coupled with the apparent inactivation of the catalytic triad across all SpmX genes suggests that muramidase function may not remain fully active.

### The SpmX muramidase domain has a wider, more dynamic catalytic cleft than related phage lysins

Obtaining the structure of the SpmX muramidase domain (residues 1-150) from *Asticcacaulis excentricus* (SpmX-Mur-*Ae*) allowed us to directly visualize the effect of the D20L and T26X mutations on the catalytic cleft. We solved the X-ray crystal structure by molecular replacement using the atomic coordinates of phage P22 lysozyme (PDB 2ANX) to 1.9 Å resolution (*R*cryst of 21.1%, *R*free of 25.5%,) (**Table S2**). SpmX-Mur-*Ae* exhibited the expected structural similarity with the phage P22 lysozyme (root mean square deviation (rmsd) of 1.7 Å and 40% identity over 141 aligned Cα atoms, Dali Z-score of 21.5). It therefore shares the characteristic structure of T4 lysozymes, with the predicted catalytic glutamate occurring at the C-terminal end of the first alpha-helix and situated in the catalytic cleft formed between the N- and C-terminal lobes (**Fig. 3A**). The active conformation of the distantly related SAR endolysin protein R21 (PDB 3HDE) from bacteriophage P21 (**Fig. S2**) is included in the structural alignment in **Figure 3A** to emphasize the ways in which the SpmX muramidase domain deviates from phage lysozymes: SpmX-Mur-*Ae* contains an extended beta-hairpin in the C-terminal lobe and, more importantly, the canonical GH beta-hairpin in the N-terminal lobe of SpmX-Mur-*Ae* splays away from the catalytic cleft relative to the GH beta-hairpins of the phage lysozymes. There is additional evidence for the flexibility of the GH beta-hairpin in the various conformations observed in the crystallography data, as among the three molecules of SpmX-Mur-*Ae* in the asymmetric unit, this region exhibited the most conformational differences. The overlay of the three chains in **Figure 3B** illustrates how the orientation of the GH beta-hairpin is tilted by about 16° between chains A and B; moreover, chain A appears to have lost some of its beta-strand secondary structure. That SpmX-Mur-*Ae* is capable of both conformations suggests a heightened flexibility in this region compared to other T4L-like lysozymes, which may reduce the ability of the enzyme to coordinate the hydrolysis of peptidoglycan in the catalytic cleft.

**Figure 3.**
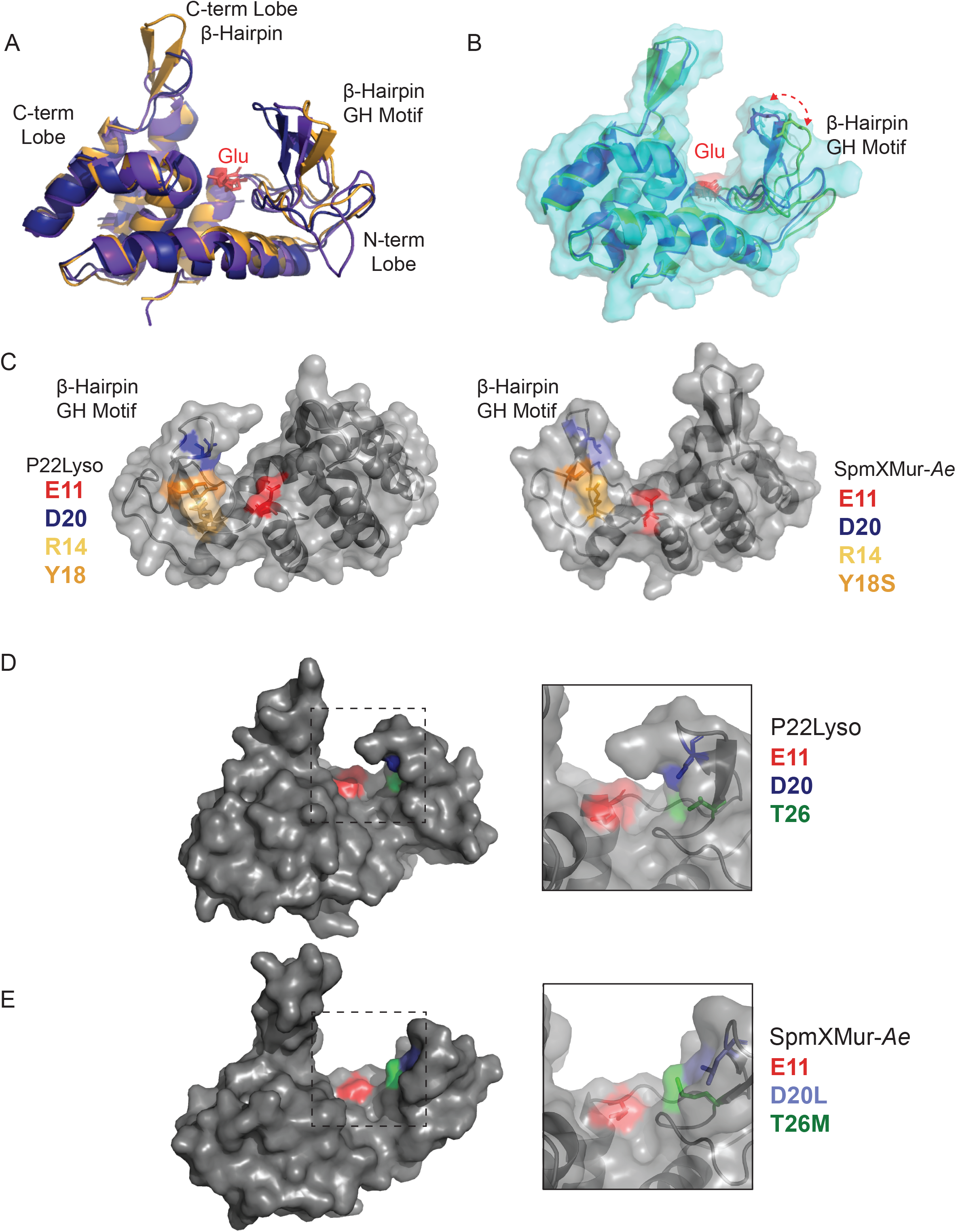
The structure of SpmX muramidase domain has a wider, more dynamic catalytic cleft than related phage lysins. **(A)** Structural alignment of P22 lysozyme (PDB 2ANX, the model used for molecular replacement) in purple, R21 endolysin from P21 (PDB 2HDE, a distantly related GH24 T4L lysozyme) in navy blue, and SpmX-Mur-*Ae* in gold (PDB 6H9D). The catalytic glutamate is shown in red. **(B)** Structural alignment of the three SpmX-Mur-*Ae* molecules, chains A (green), B (light blue), and C (dark blue), from the asymmetric unit. The surface of chain B is shown in partially transparent light blue. The double-headed arrow indicates the tilt of about 16° between the GH beta-hairpins of chains B and A. **(C)** Overlays of ribbon diagrams and surfaces of P22 lysozyme (2ANX, left) and SpmX-Mur-*Ae* (6H9D, right) illustrating the conformation of the critical residues E11 (red), D20 (dark blue), R14 (yellow), and Y18 (orange). T4L numbering is used for ease of comparison. These structures have been rotated 180° around the y axis from their representation in (A, B, D, E)**. (D)** Surface representation of P22 lysozyme (2ANX) with inset showing ribbon diagram and conformation of catalytic cleft with the canonical E11/D20/T26 catalytic triad. **(E)** Surface representation of SpmX-Mur-*Ae* (6H9D) with inset showing ribbon diagram and conformation of remodeled catalytic cleft with E11/D20L/T26M.

Sequence comparisons of the GH motif in **Figure 2B** show that SpmX muramidase domains have lost a highly conserved tyrosine residue at position 18. Although T4L enzymatic activity is not sensitive to mutation at this position [27], it is invariant across all the phage lysozyme classes we analyzed. Visualization of Y18 in the P22 lysozyme structure (**Fig. 3C**) shows that it interacts with R14 at the base of the beta-hairpin, possibly a critical interaction for coordinating the beta-hairpin with the catalytic glutamate. In SpmX-Mur-*Ae*, Y18S still appears to make hydrogen-bonding contact with R14; however, most SpmX muramidase domains have non-polar residues at position 18 (**Fig. 2B(iv)),** which may reduce coordination between the catalytic glutamate and the GH beta hairpin. It has been previously shown that the Y18 position is a hot-spot for compensatory mutations that restore activity to inactive catalytic mutants [31], and it is intriguing to imagine that mutations at this position in SpmX muramidase are the result of a remodeled catalytic cleft.

**Figure 3** shows the catalytic clefts of both P22 lysozyme (**D**) and SpmX-Mur-*Ae* (**E**). In P22 lysozyme, the E11 carbonyl, D20 carboxyl, and T26 hydroxyl groups point into the aqueous catalytic cleft. In SpmX-Mur-*Ae,* the cleft is slightly reorganized, with the S-methyl thioether of T26M still within 20 Å of E11 and potentially capable of interacting with peptidoglycan. In about two thirds of the SpmX genes, position 26 has either a valine or an isoleucine, which have no polar moieties to contribute to the cleft (**Fig. 2B(iv)**). With the structural data from SpmX-Mur-*Ae*, we can infer that the SpmX muramidase domain has a remodeled catalytic cleft with a correctly positioned catalytic glutamate. However, the increased flexibility between the GH motif and the glutamate, as well as the loss of key coordinating residues might reduce, if not eliminate, hydrolytic activity of SpmX.

### SpmX retains reduced hydrolytic activity on peptidoglycan

To determine if the SpmX muramidase domain was capable of binding peptidoglycan, various constructs from *C. crescentus, A. excentricus,* and *A. biprosthecum* were purified and incubated with sacculi isolated from the three species. Both muramidase domains and entire soluble domains (muramidase with intermediate domain) from all three species, including the E11A and N105R mutants of SpmX from *C. crescentus* bound sacculi from all three species (**Fig. S3**). Despite previous reports of inactivity [18], we found that purified SpmX muramidase exhibited hydrolytic activity. We used remazol brilliant blue (RBB) assays, which are standard for measuring lysozyme activity [32], to compare the activity of SpmX muramidase from *C. crescentus* (SpmX-Mur-*Cc*) to that of the P22 lysozyme (P22Lyso) and a P22Lyso mutant with a similarly inactivated catalytic cleft (P22Lyso-D20L) (**Fig. 4A**). The activity curves demonstrate that both SpmX-Mur-*Cc* and P22Lyso-D20L exhibit similarly attenuated hydrolytic activity in comparison to P22Lyso: both reached maximal levels of RBB release near enzyme concentrations of 15 μM while P22Lyso reached the same levels near 5 μM. Mutants in which the catalytic glutamate was replaced with alanine (SpmX-Mur-*Cc-*E11A and P22Lyso-E11A) did not exhibit activity (**Fig. S4A**). Restoring the ancestral D20 (SpmX-Mur-*Cc-*L20D) did not increase SpmX activity *in vitro* (**Fig. S4B**). These data indicate that the “inactivating” substitution D20L attenuates enzymatic activity whereas mutating the catalytic glutamate abolishes it altogether.

**Figure 4.**
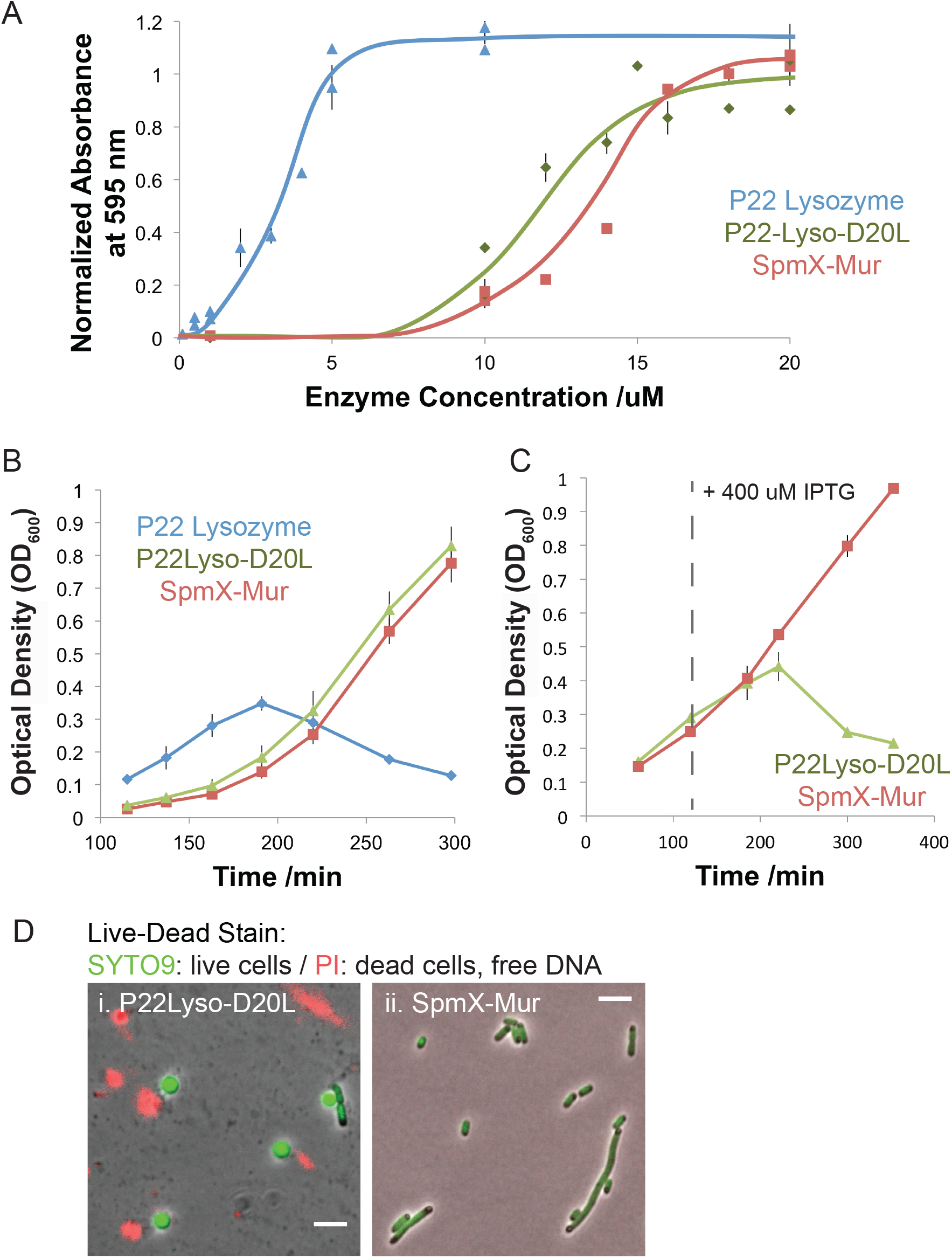
The D20L mutation attenuates P22 hydrolytic activity. **(A)** Remazol brilliant blue assays on *C. crescentus* sacculi using purified P22 lysozyme, P22 lysozyme D20L mutant, and *C. crescentus* SpmX muramidase. Active enzymes release peptidoglycan monomers covalently-bound to RBB into the supernatant that are detected by absorbance at 595 nm. Error bars are ± standard deviation. Lines are drawn to help guide the eye toward basic trends. Data points are from various days and sacculi preparations, but with internal normalization to Hen Egg White Lysozyme (HEWL). **(B** and **C)** Growth curves of Lemo21(DE3) *E. coli* expressing P22 lysozyme (blue), P22 lysozyme D20L mutant (green), and *C. crescentus* SpmX muramidase (red). Proteins were expressed from pET22b with a N-terminal PelB signal sequence. In **(B)**, strains were grown in 5 mM rhamnose without IPTG for maximal repression of basal expression from the plasmids. In **(C)**, strains were grown without rhamnose and induced with 400 μM IPTG at the indicated time. **(D)** Phase/fluorescent overlays show live/dead staining of (i) Lemo21(DE3) cells expressing P22Lyso-D20L and (ii) SpmX-Mur-*Cc* after four hours of induction. Green, membrane permeable SYTO 9 stains DNA in live cells and red, membrane impermeable propidium iodide nucleic acid dyes labels released nucleoids and DNA from lysed bacteria. The rounding of the *E. coli* in (i) is characteristic of spheroplast formation and lysis by hydrolytic activity on the cell wall. Scale bars are 5 μm.

We were surprised by the activity of P22Lyso-D20L and SpmX-Mur given the reported inhibition of lytic activity of T4L with D20 mutations in phage plaque assays [27]. We were interested in the biological implications of this *in vitro* activity, and whether the attenuated mutant was active in the periplasmic environment. We therefore designed an experimental system to test the activity of P22Lyso and SpmX-Mur-*Cc* mutants in the periplasmic space of *E. coli*. In this system, we expressed the various muramidase constructs with N-terminal PelB leader sequences (pET22b) for periplasmic expression in Lemo21(DE3) cells, which allow tunable expression of toxic products (see Methods). Lemo21(DE3) cells carrying P22Lyso lysed even in the absence of induction (**Fig. 4B)**. In contrast, Lemo21(D32) cells carrying P22Lyso-D20L lysed only after induction, and strains carrying SpmX-Mur-*Cc* and P22Lyso-E11A never lysed (**Fig. 4C** and **S4C**) despite equivalent periplasmic expression levels (**Fig. S4D**). Different growth conditions and media did not affect the viability of cells expressing SpmX-Mur-*Cc* (**Fig. S4C**). Moreover, SpmX-Mur-*Cc* was active on sacculi isolated from Lemo21(DE3), eliminating the possibility that it cannot cleave *E. coli* peptidoglycan (**Fig. S4E**). It remains unclear why the periplasmic activity of SpmX-Mur-*Cc* does not match its *in vitro* activity, as it appears to have a similar activity to P22Lyso-D20L and yet surpasses the periplasmic levels at which P22Lyso-D20L lyses *E. coli*. It is possible that SpmX-Mur has additional mutations that either make it difficult to fold correctly in the *E. coli* periplasm, or inactivate its activity in the periplasmic environment. However, the periplasmic expression tests in *E. coli* confirm that the D20L mutation attenuates the hydrolytic activity of P22 lysozyme. This tuning of enzymatic activity might have served as a critical detoxifying step in the co-option of the muramidase domain from phage. Moreover, such attenuation helps explain why the D20L substitution is absent from phage despite broad conservation among *spmX* alleles. Because SpmX has retained the ancestral catalytic glutamate and its modified catalytic cleft is capable of hydrolytic activity, we conclude that this attenuated activity is under purifying selection in SpmX and important for SpmX function.

### Inactivating the muramidase domain impacts SpmX localization *in vivo*

To investigate the role of the conserved catalytic glutamate in SpmX function *in vivo*, chromosomal E11A mutants (E19A in SpmX numbering) were made in *C. crescentus*, *A. excentricus*, and *A. biprosthecum* with eGFP gene fusions (**Fig. 5**). We determined the effects of the E11A mutation on cellular morphology, as the Δ*spmX* strains in all three species have morphological phenotypes (**Fig. 5ABCii**). In *C. crescentus*, Δ*spmX* cells have a characteristic elongated morphology resulting from failed division cycles and often grow stalks prematurely from daughter cells that fail to divide completely (**Fig. 5Aii**) [15]. In *Asticaccaulis*, ∆*spmX* cells lack stalks and do not appear to have other developmental phenotypes (**Fig. 5BCii**) [17]. In *C. crescentus*, the E11A mutant population contained both WT-like cells and cells exhibiting the division defect, but with less severity than in ∆*spmX* (**Fig. 5Aiii**). In both *Asticcacaulis* species, the E11A mutants still grew stalks (**Fig. 5BCiii**). Nevertheless, the *A. biprosthecum* E11A mutant exhibited a significant loss of bilateral stalks (3.5 fold reduction) and an increase in the frequency of cells with a single stalk (**Fig. S5D**).

**Figure 5.**
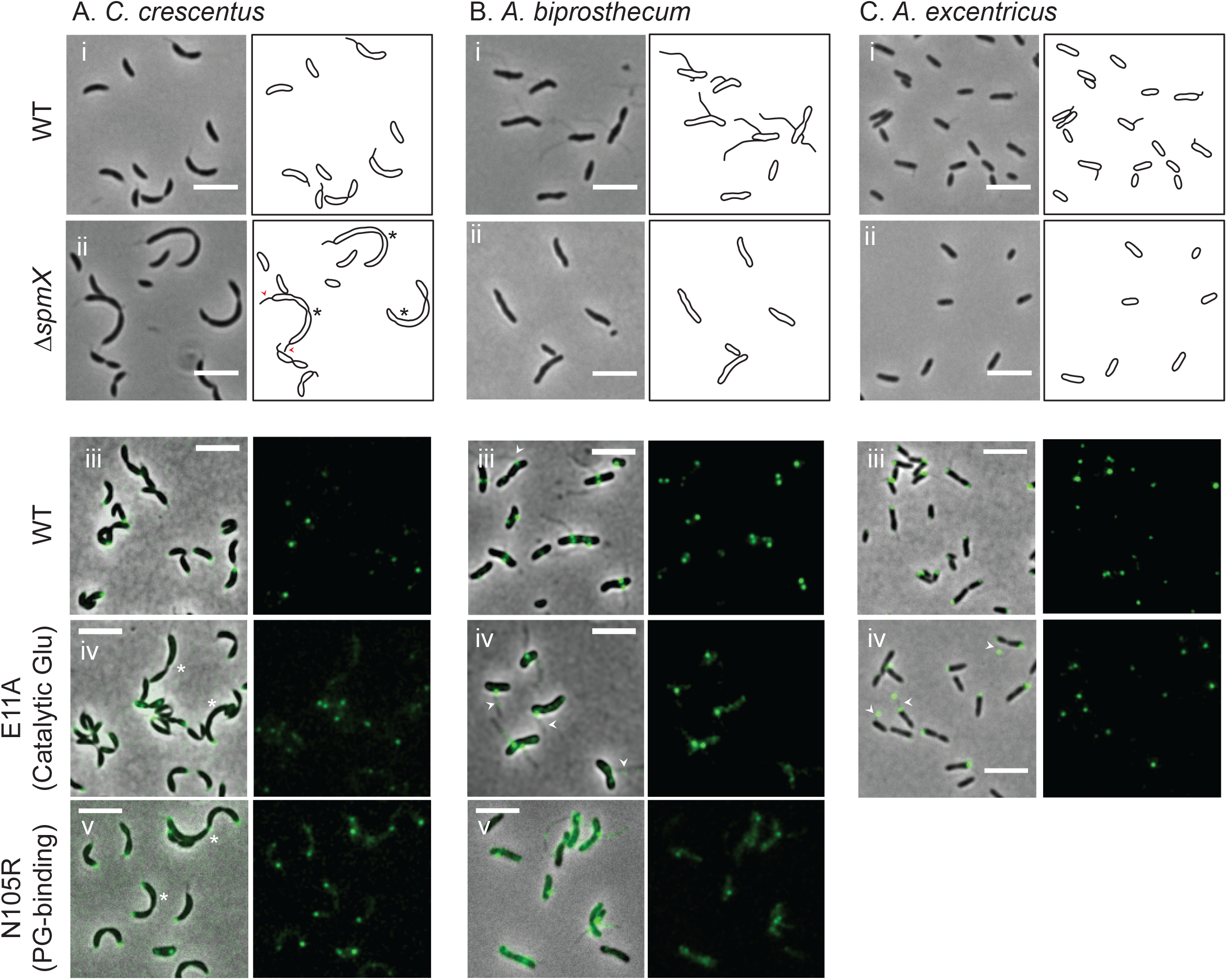
Inactivating the muramidase domain partially delocalizes SpmX *in vivo*. Phase and fluorescent images of (**A)** *C. crescentus*, (**B)** A*. biprosthecum,* and **(C)** *A. excentricus.* In the top panel, phase images with derived schematics emphasizing stalks and morphologies are shown for (**i**) WT and (**ii**) ∆*spmX* cells. In **Aii**, *C. crescentus* cells exhibiting characteristic Δ*spmX* divisional defects are marked with asterisks and a cell growing stalks from both poles has its stalks marked with red arrowheads. Phase and fluorescent images of cells expressing (**iii**) SpmX-eGFP, (**iv**) SpmX-E11A-eGFP, or (**v)** SpmX-N105R-eGFP from the native chromosomal locus are shown in the lower panels. In **Aiv** and **Av**, cells with divisional defects are marked with white asterisks. In **Biii** and **Biv**, cells with one lateral or subpolar stalk are marked with white arrowheads. In **Civ**, cells with foci at the tips of stalks are marked with white arrowheads. All scale bars are 5 μm.

We used GFP fusions of WT and mutant SpmX to monitor changes in SpmX localization. In all species, WT SpmX localizes at the future position of the stalk, either at the pole as in *C. crescentus*, or at sub-polar or bilateral positions in *Asticcacalis*, and is retained at this position (**Fig. 5ABCiii**). Both *C. crescentus* and *A. biprosthecum spmX* E11A mutants exhibited an increase in mislocalized SpmX throughout the cell body compared to WT (**Fig. 5ABiv**). Quantification of the fluorescence data indicated that while the overall mean fluorescence of the cells were the same as WT, the SpmX foci were significantly less intense in the mutants (**Fig. S5AB**). We also observed a 3X increase in SpmX E11A in the stalks of *A. biprosthecum* compared to WT (**Fig. S5B**). Although no difference in focal fluorescence intensity was observed in *A. excentricus spmX* E11A mutant cells, more cells had a second SpmX focus at the tip of the stalk than WT cells (**Fig. 5iv, S5C)**, indicating altered localization. Together these data show that the E11A mutation impacts SpmX localization in all three species.

N/Q105 has been shown to coordinate peptidoglycan in the active cleft of T4L [33] and the mutation Q105R abolished activity in T4 phage plaque assays [27]. The mutation N105R (N91R in SpmX numbering) in *C. crescentus* and *A. biprosthecum* resulted in similar delocalization phenotypes as E11A (**Fig. 5BCiiv**). Western blots of cells expressing WT SpmX-eGFP and SpmX mutants indicated that the delocalization was not due to clipping (**Fig. S5E**). Overall, these data show that eliminating the catalytic glutamate affects SpmX localization and function. Although it is not clear whether the E11A mutation specifically affects the potential hydrolytic activity or the peptidoglycan-binding ability, the similar phenotype from mutating a predicted peptidoglycan-interacting residue (N105R) underscores the importance of SpmX-peptidoglycan interactions. That the catalytic mutant has an intermediate morphological phenotype in *C. crescentus* and one *Asticcacaulis* species indicates that the muramidase domain may coordinate SpmX functions similarly in the two genera and that this function may rely on its enzymatic activity.

### Replacing the muramidase domain, or removing it, interferes with native SpmX protein levels *in vivo*

In an effort to understand the role of the highly conserved muramidase fold in SpmX function, we made chimeras wherein P22 lysozyme replaced the muramidase domain of SpmX in *C. crescentus* (P22Lyso-SpmX). While similarly made constructs with SpmX and the SpmX-E11A mutant exhibited the previously determined morphological and delocalization phenotypes (**Fig. 6Aii-iii**), replacing the SpmX muramidase domain with P22 lysozyme phenocopied the parent ∆*spmX* strain. However, we were unable to detect any sfGFP-fusion products in these chimeras by Western blot (**Fig. 6B**). We confirmed that P22Lyso-SpmX had been correctly inserted in these strains by sequencing, suggesting that the chimeras were expressed but degraded quickly in *C. crescentus*. Therefore the phenotype of this chimera is due to loss of SpmX and not the addition of the P22 lysozyme domain. Inactivation of P22Lyso (E11A) did not change the outcome, suggesting that the toxicity of the phage muramidase was not driving degradation of SpmX. Deletion of the muramidase domain from the *spmX* locus in all three species also resulted in strains that failed to produce detectable amounts of Δmur-SpmX-sfGFP by Western blot (**Fig. S5E**). These results suggest that the SpmX muramidase domain is necessary to produce and/or maintain WT levels of SpmX in all three species, and that phage muramidase P22Lyso, with high sequence similarity (51%) and structural homology (RMSD 1.7 Å), is not sufficient to replace it.

**Figure 6.**
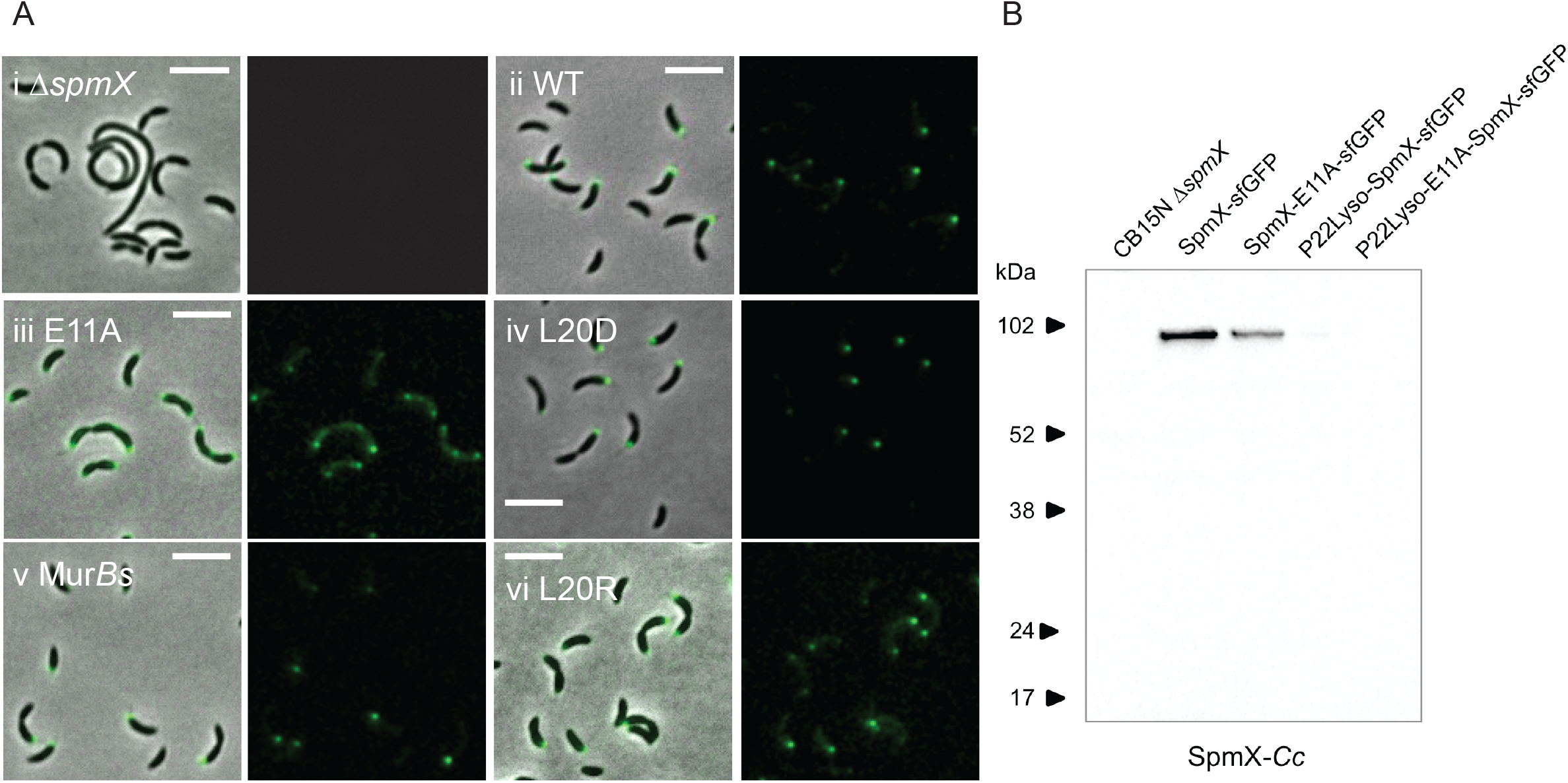
Replacing the muramidase domain interferes with native SpmX protein levels *in vivo*. **(A)** Phase and fluorescent images of strains in which the native *spmX* allele was replaced with the following gene fusions in the Δ*spmX* parent strain (i): (ii) WT *spmX-sfGFP*, (iii) *spmX-E11A-sfGFP*, (iv) *spmX-L20D-sfGFP*, (v) *MurBs-*Δ*mur-SpmX-sfGFP* where *MurBs* is the muramidase domain from *Brevundimonas subvibrioides* SpmX, and (vi) *spmX-L20R-sfGFP*. All scale bars are 5 μm.. **(B)** Western blot comparing the ∆*spmX* parent strain to SpmX mutants and chimeras inserted at the *spmX* locus. In all cases, the primary antibody is directed against the C-terminal GFP fusion.

Given that P22Lyso and SpmX muramidase are fairly distant relatives despite high sequence conservation, we tested the ability of other SpmX muramidase domains to replace that of *C. crescentus*. First, we attempted to restore the phage active site D20 (L28D in SpmX numbering) in the native SpmX gene, but this had no evident effect on SpmX localization or cell morphology (**Fig. 6iv**). Previously, the muramidase domains of *C. crescentus* and *Asticcacaulis* were shown to be interchangeable [17], so we extended the sequence distance to SpmX muramidases from the next closest relative *B. subvibrioides*, which has D20R in the catalytic cleft, and the most distant relative *P. bermudensis,* which shares the D20L mutation. We found that the muramidase domain from *B. subvibrioides* supported the WT phenotype in *C. crescentus* (**Fig. 6v)**, but that the SpmX muramidase domain from *P. bermudensis* did not. We were surprised to see no evidence of delocalization in the *B. subvibrioides* SpmX chimera because the L20R mutant of SpmX in *C. crescentus* showed some delocalization (**Fig. 6vi**). This result suggests that the L20R mutation in the brevundimonads must coexist with other compensatory mutations. The SpmX muramidase domain from *P. bermudensis*, like P22Lyso, must be too distant from *C. crescentus* to support WT levels of SpmX protein. In combination with data from the P22Lyso chimeras, these data indicate that a T4L GH fold alone is not sufficient for SpmX function, and that the SpmX muramidase domain must contain mutations necessary for stable protein levels in Caulobacterales. It also suggests that this domain has additional constraints on it unrelated to potential peptidoglycan interactions.

## Discussion

Bacteriophages shape bacterial evolution in various ways: they increase bacterial diversity by selectively preying on species [2,34], drive horizontal gene transfer of phage and bacterial genes [8,35], and serve as reservoirs of raw material for new bacterial genes [36,37]. Phages are heralded as a major source of genetic material for novel gene emergence in bacteria [2,36,37], but, as we discuss later in this section, very few examples of novel gene emergence from prophage exist in the literature. We have investigated the origin and function of a taxonomically restricted gene from Caulobacterales, *spmX*, and determined that its occurrence is the result of the fusion and domestication of a peptidoglycan hydrolase gene commonly found in prophage. Despite the previously demonstrated role of SpmX as a scaffold in developmental regulation and morphology, the muramidase domain retains high sequence similarity to phage lysozymes, which are toxic to bacteria. The active cleft contains mutations that have attenuated the toxic activity of the domain, presumably making it available for genetic innovation and bacterial use. We show here that the domain remains enzymatically active on peptidoglycan and that reducing or eliminating this activity alters the function of the full-length protein *in vivo*. Thus, the SpmX gene represents a bacterial gene innovation, specific to the Caulobacterales order and with housekeeping function, that originally arose from a prophage gene with antibacterial activity.

Previously, it was suggested that the SpmX muramidase domain functions only in protein-protein and self-oligomerizing interactions in SpmX’s role as a developmental regulator and scaffold in *C. crescentus* [18]. This conclusion was based on the lack of detectable activity from the purified domain and the inability of the catalytic E11R (E19R in SpmX numbering) mutant to self-oligomerize. The conservation of the GH fold and the catalytic glutamate clearly suggests that the activity of SpmX is important for its domesticated function, and, importantly, we were able to observe muramidase activity *in vitro* with purified protein. It is highly likely that the E11R mutation greatly destabilizes the structure of the muramidase domain. We found that the E11A mutant eluted in multiple fractions during purification, indicating decreased conformational stability. Moreover, the E11R mutant exactly phenocopies the Δmur-SpmX phenotype *in vivo* in that the protein product was no longer detectable in the cells expressing the gene [18]. In this sense, the effect of the E11R mutation is similar to using a distantly related muramidase domain (like P22Lyso) or deleting portions of the muramidase domain entirely. These data indicate that the muramidase domain plays an unanticipated role in maintaining stable levels of SpmX protein across all tested species: without the muramidase domain, the protein is misfolded or misprocessed and is therefore quickly degraded.

Inactivating the SpmX muramidase domain resulted in developmental defects in *C. crescentus* and significant loss of the bilateral stalk morphology in *A. biprosthecum*. It is curious that inactivating the enzymatic domain did not yield a null phenotype or complete delocalization. Since SpmX-E11A and N105R still bind to sacculi *in vitro*, it is possible that enough peptidoglycan-interactions are maintained in the mutants for partial SpmX function. It is also possible that SpmX recruits another protein with redundant enzymatic activity that cannot be recruited in the Δ*spmX* mutant, as it is already known that SpmX interacts with targeting factors via its C-terminal domains in *Asticcacaulis* [17] and possibly via its transmembrane segments with DivJ [15]. Finally, it is hard to distinguish whether there is a direct relationship between catalytic activity and peptidoglycan binding, and therefore localization, or if cleavage of peptidoglycan by SpmX could indirectly impact SpmX localization. The multiple domains and pleiotropic effects of SpmX make it difficult to assess the effects of an individual domain on its *in vivo* function. However, our data support a model in which the muramidase domain of SpmX is still active, and this activity is used to localize SpmX directly or indirectly.

SpmX emerges in the genomic record at the root of Caulobacterales with the attenuating D20L mutation already in the muramidase domain (**Fig. 1B**). The D20L mutation is therefore ancestral and potentially the initial step in the co-option of the domain. The conservation of D20L throughout most of Caulobacterales suggests evolutionary constraint on this position despite no observable phenotype from the SpmX-Mur-*Cc* L20D reversion mutation *in vivo* or *in vitro*. There are several modifications to the active cleft, including the loss of selection on the third catalytic triad position, T26, and the invariant residue Y18. This pair is interesting in that Y18 was identified as a hotspot for spontaneous second site revertants of T26 mutants in T4L [31]. It is possible that the scatter we see at these two positions are compensatory mutations retaining attenuated activity, although a clear history of covariation is not clear. In two groups, SpmX has diverged from the ancestral D20L mutation: the marine genera *Oceanicaulis* and *Maricaulis* (D20R/G), and the freshwater/soil genus *Brevundimonas* (D20R) (**Fig. 1B, S1**). Interestingly, D20R is covariant with residue N105 (**Fig. S1**), which is a peptidoglycan-interacting residue in T4 lysozyme [33]. The covariance of peptidoglycan-interacting residues in these diverging genera further underscores the importance of this domain in peptidoglycan interactions, rather than just protein-protein interactions.

The SpmX gene has arisen recently enough to see the hallmarks of novel gene emergence and adaptation in a constrained bacterial clade. The gene either arose from a fusion event in the bacterial genome, or the original phage gene contained the transmembrane segments. Domestication of the muramidase appears concomitant with the occurrence of the fused SpmX domains in the extant genetic record. Maintenance of the muramidase domain since the emergence of SpmX and its activity in current living members of Caulobacterales suggest that its activity was selected for in the ancestral protein and still implicated in its modern function. Given the variability of the intermediate domain throughout Caulobacterales (**Fig. 1A**), this domain is under few constraints and appears to have undergone multiple independent events of elaboration and reduction in this order. This region of charged residues and prolines drives SpmX self-oligomerization *in vitro* [18], and may also facilitate other protein interactions. For example, the intermediate domain is responsible for the targeting of SpmX to sub-polar and bilateral positions in *Asticcacaulis* [17].

In several reported cases bacteria have domesticated phage genes for genetic manipulation and transfer, bacterial warfare, virulence, and secretion. However, these events are distinct from that which created the novel bacterial gene SpmX. Phage genes for DNA replication and recombination have replaced bacterial functional homologues within bacterial genomes several times [38–41]. These genes retain their original function and are now used by bacteria to carry out the same tasks. Gene transfer agents (GTAs) pose an interesting case where structural proteins from domesticated cryptic prophage are used by bacteria to package random DNA from the bacterial genome to presumably share with other bacteria [42]. In Alphaproteobacteria, a specific GTA has been stably maintained across several bacterial orders but the beneficial function is not precisely known [43,44]. Regardless, this domesticated island of phage genes still shuttles DNA around, as it once did in ancestral infectious cycles. Phage tails appear to have been weaponized many times, with multiple domestication events resulting in type VI secretion systems [45,46], tailocins and phage tail-like bacteriocins [12–14], phage tail-like systems with insecticidal properties [11,47,48], and phage tail-like arrays [49]. All of these represent a “guns for hire” acquisition scheme in which phage genes are co-opted without loss of ancestral toxicity and function [2]. Many of these genes reside in genomic islands and confer environmental, niche-specific advantages that directly exploit their ancestral activity for the benefit of the host. Similarly, in two other known cases of phage lysozyme domestication in bacteria, structurally similar, active domains have been fused to colicins [50] or are predicted to be secreted with type III secretion systems [51], presumably for use in bacterial warfare or infection. In one strange case, a phage lysozyme gene has been co-opted in bivalve genomes, which apparently still use the gene for its antibacterial properties [52].

The domestication of the muramidase domain in SpmX is distinct from the above cases of “guns for hire” because the phage gene has been incorporated into a novel bacterial gene with new function in basic cellular processes. The SpmX muramidase domain, although active, is no longer used to lyse bacterial cells. Instead, the active domain plays a role in localizing the SpmX protein for its function in developmental regulation and morphogenesis and has become part of a core gene in Caulobacterales. The co-option of phage genes for housekeeping function is likely a common event in nature, but identifying such genes may require a careful search. We suggest that the characteristics of SpmX may recommend future strategies for their detection: searching for phage gene homologues that have long histories of vertical inheritance and signs of bacterial innovation.

## Acknowledgements

We thank members of the Brun, Vernet, and Dessen laboratories for support, advice and encouragement. Many thanks to members of the Brun Laboratory, Ernesto Vargas, Farrah Bashey-Visser, Jay Lennon, Daniel Schwartz, and Breah LaSarre for critical reading and editing of the manuscript. Additional thanks to critical preprint review by the journal clubs of the (Pamela) Brown Lab at the University of Missouri and the Süel Lab at UCSD. We thank David Flot (ESRF, beamline ID30a1) for support in data collection. Support for this work came from NIH Grant 2R01GM051986 and R35GM122556 (to Y.V.B.), and NIH National Research Service Award F32GM112362 (to A.M.R.). Work of Y.V.B. at the Institut de Biologie Structurale in Grenoble was supported by a Fulbright US Research Scholar Award. This work used the platforms of the Grenoble Instruct-ERIC Center (ISBG : UMS 3518 CNRS-CEA-UGA-EMBL) with support from FRISBI (ANR-10-INSB-05-02) and GRAL (ANR-10-LABX-49-01) within the Grenoble Partnership for Structural Biology (PSB).

---

**Table S1.** *spmX* genes

| Gene ID | Genome ID | Genome Name | Aas |
| --- | --- | --- | --- |
| 2579190773 | 2576861677 | *Asticcacaulis benevestitus* DSM 16100, ATCC BAA-896 | 825 |
| 2515453797 | 2515154077 | *Asticcacaulis benevestitus* DSM 16100, ATCC BAA-896 | 825 |
| 651318754 | 651285000 | *Asticcacaulis biprosthicum* C19, ATCC 27554 | 826 |
| 649817473 | 649633007 | *Asticcacaulis excentricus* CB 48 | 810 |
| 2579161412 | 2576861669 | *Asticcacaulis* sp. AC402 | 861 |
| 2575314089 | 2574180013 | *Asticcacaulis* sp. AC460 | 803 |
| 2573897243 | 2571042840 | *Asticcacaulis* sp. AC466 | 863 |
| 2579118533 | 2576861656 | *Asticcacaulis* sp. YBE204 | 851 |
| 2597242146 | 2596583650 | *Asticcacaulis taihuensis* CGMCC 1.3431 | 775 |
| 2525860150 | 2524614798 | *Brevundimonas aveniformis* DSM 17977 | 453 |
| 2562052386 | 2561511096 | *Brevundimonas bacteroides* DSM 4726 | 507 |
| 2534536731 | 2531839707 | *Brevundimonas diminuta* 470-4 | 291 |
| 2635997287 | 2634166514 | *Brevundimonas diminuta* XGC1 | 291 |
| 2580800557 | 2579778742 | *Brevundimonas naejangsanensis* B1 | 291 |
| 2525302517 | 2524614659 | *Brevundimonas naejangsanensis* DSM 23858 | 296 |
| 2602030628 | 2600255394 | *Brevundimonas nasdae* TPW30 | 393 |
| 2597236398 | 2596583649 | *Brevundimonas* sp. 374 | 381 |
| 2646174451 | 2645727605 | *Brevundimonas* sp. AAP58 | 493 |
| 647574062 | 647533115 | *Brevundimonas* sp. BAL3 | 526 |
| 2628481828 | 2627853674 | *Brevundimonas* sp. KM4 | 381 |
| 648120868 | 648028010 | *Brevundimonas subvibrioides* ATCC 15264 | 514 |
| 2671799930 | 2671180176 | *Brevundimonas viscosa* CGMCC 1.10683 | 420 |
| 637088217 | 637000061 | *Caulobacter crescentus* CB15 | 431 |
| 643620503 | 643348526 | *Caulobacter crescentus* NA1000 | 431 |
| 2522234411 | 2522125057 | *Caulobacter crescentus* OR37 | 429 |
| 2585155721 | 2582581280 | *Caulobacter henricii* CF287 | 419 |
| 2585146921 | 2582581279 | *Caulobacter henricii* OK261 | 398 |
| 2585198327 | 2582581293 | *Caulobacter henricii* YR570 | 419 |
| 646754844 | 646564519 | *Caulobacter segnis* ATCC 21756 | 413 |
| 2511123337 | 2510917020 | *Caulobacter* sp. AP07 | 425 |
| 2528897460 | 2528768059 | *Caulobacter* sp. JGI 0001010-J14 | 422 |
| 2528898539 | 2528768060 | *Caulobacter* sp. JGI 0001013-D04 | 421 |
| 641563661 | 641522612 | *Caulobacter* sp. K31 | 417 |
| 2587919405 | 2585428106 | *Caulobacter* sp. OV484 | 439 |
| 2686817300 | 2684623106 | *Caulobacter* sp. UKL13 | 439 |
| 2593324358 | 2590828858 | *Caulobacter* sp. UNC279MFTsu5.1 | 421 |
| 2565991034 | 2565956508 | *Caulobacter sp. UNC358MFTsu5.1* | 424 |
| 2522923390 | 2522572103 | *Caulobacter sp. URHA0033* | 413 |
| 2677566563 | 2675903340 | *Caulobacter vibrioides T5M6* | 426 |
| 2525553380 | 2524614726 | *Hellea balneolensis DSM 19091* | 439 |
| 2519259833 | 2519103046 | *Henriciella marina DSM 19595* | 535 |
| 644903978 | 644736375 | *Hirschia baltica ATCC 49814* | 597 |
| 2520004565 | 2519899518 | *Hirschia maritima DSM 19733* | 577 |
| 2578549711 | 2576861496 | *Hyphomonas adhaerens MHS-3* | 386 |
| 2578539236 | 2576861493 | *Hyphomonas hirschiana VP5* | 421 |
| 2578553211 | 2576861497 | *Hyphomonas jannaschiana VP2* | 385 |
| 2578549047 | 2576861495 | *Hyphomonas johnsonii MHS-2* | 409 |
| 638143841 | 637000135 | *Hyphomonas neptunium ATCC 15444* | 421 |
| 2578534396 | 2576861492 | *Hyphomonas oceanitis SCH89* | 407 |
| 2578543740 | 2576861494 | *Hyphomonas polymorpha PS728* | 418 |
| 2583291548 | 2582580773 | *Hyphomonas sp. 22III-22F38* | 390 |
| 2583281302 | 2582580770 | *Hyphomonas sp. 25B14_1* | 392 |
| 2583297317 | 2582580774 | *Hyphomonas sp. BH-BN04-4* | 407 |
| 2583284155 | 2582580771 | *Hyphomonas sp. CY54-11-8* | 386 |
| 2583256535 | 2582580764 | *Hyphomonas sp. L-53-1-40* | 398 |
| 2583287429 | 2582580772 | *Hyphomonas sp. T16B2* | 392 |
| 638141228 | 637000157 | *Maricaulis maris MCS10* | 638 |
| 2635165820 | 2634166316 | *Maricaulis salignorans DSM 16077* | 633 |
| 2554749235 | 2554235119 | *Maricaulis sp. JL2009/Euryhalocaulis caribicus* | 464 |
| 2523851441 | 2523533582 | *Oceanicaulis alexandrii DSM 11625* | 643 |
| 638892666 | 638341138 | *Oceanicaulis alexandrii HTCC2633* | 596 |
| 2609050000 | 2608642188 | *Oceanicaulis sp.* | 514 |
| 2582569245 | 2579779170 | *Oceanicaulis sp. HL-87* | 514 |
| 648155025 | 648028050 | *Parvularcula bermudensis HTCC2503* | 344 |
| 2574131646 | 2571042909 | *Phenylobacterium composti DSM 19425* | 438 |
| 642759179 | 642555147 | *Phenylobacterium zucineum HLK1* | 445 |
| 2524841340 | 2524614539 | *Ponticaulis koreensis DSM 19734* | 687 |
| 2522300904 | 2522125083 | *Robiginitomaculum antarcticum DSM 21748* | 502 |
| 2522254887 | 2522125070 | *Woodsholea maritima DSM 17123* | 853 |

**Table S2.**
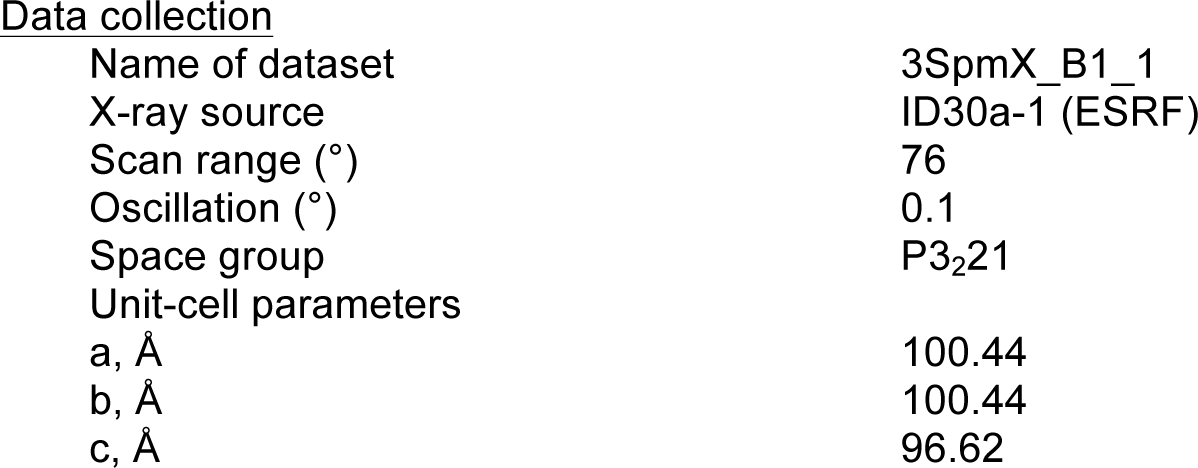
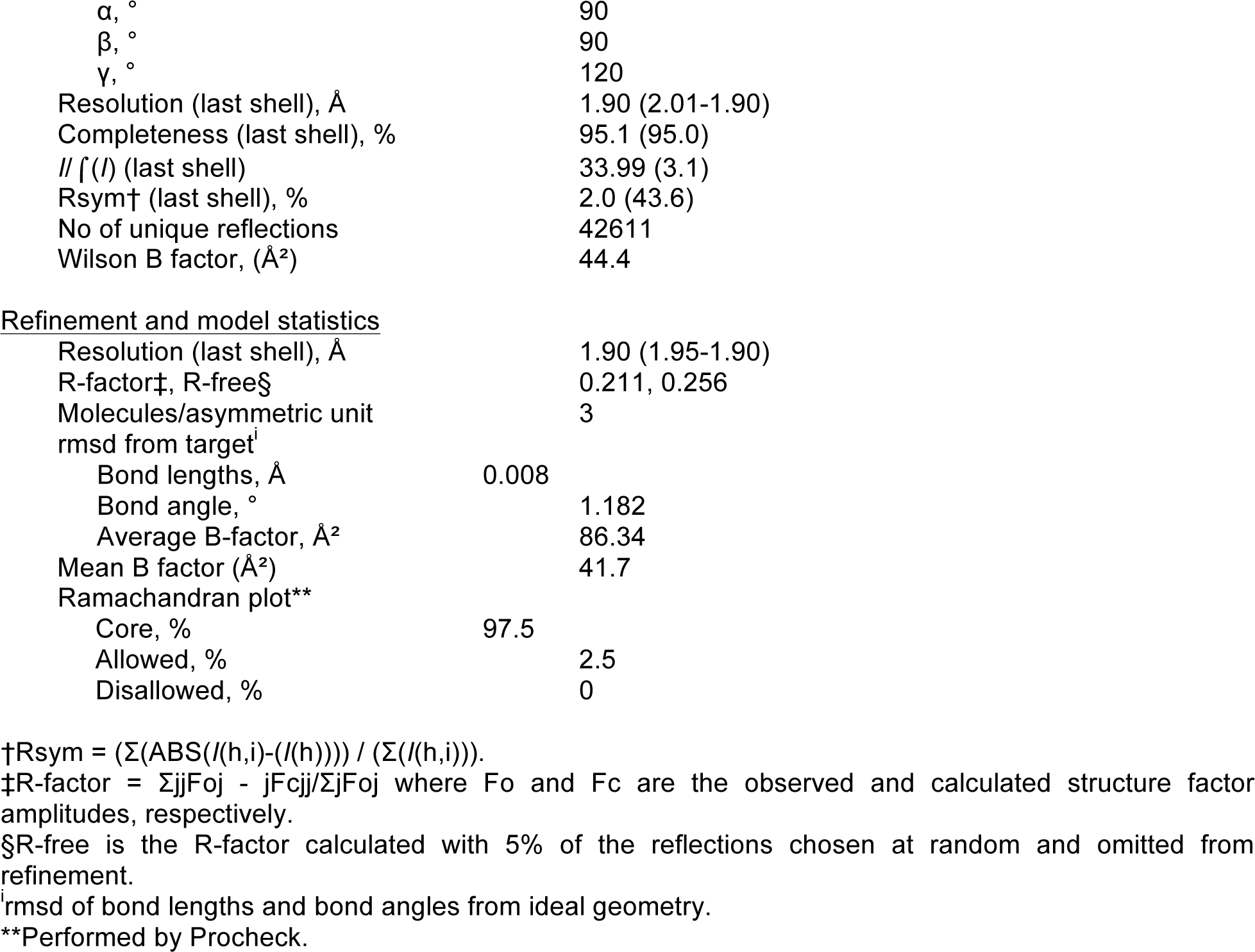
Data collection and refinement statistics

|  |  |
| --- | --- |
| Name of dataset | 3SpmX_B1_1 |
| X-ray source | ID30a-1 (ESRF) |
| Scan range (°) | 76 |
| Oscillation (°) | 0.1 |
| Space group | P3<sub>2</sub>21 |
| Unit-cell parameters |  |
| a, Å | 100.44 |
| b, Å | 100.44 |
| c, Å | 96.62 |

|  |  |  |
| --- | --- | --- |
| 570 | $\alpha$, ° | 90 |
| 571 | $\beta$, ° | 90 |
| 572 | $\gamma$, ° | 120 |
| 573 | Resolution (last shell), Å | 1.90 (2.01-1.90) |
| 574 | Completeness (last shell), % | 95.1 (95.0) |
| 575 | $\ I(I)\ $ (last shell) | 33.99 (3.1) |
| 576 | Rsym† (last shell), % | 2.0 (43.6) |
| 577 | No of unique reflections | 42611 |
| 578 | Wilson B factor, (Å<sup>2</sup>) | 44.4 |

Refinement and model statistics
|  |  |  |
| --- | --- | --- |
| 581 | Resolution (last shell), Å | 1.90 (1.95-1.90) |
| 582 | R-factor‡, R-free§ | 0.211, 0.256 |
| 583 | Molecules/asymmetric unit | 3 |
| 584 | rmsd from target<sup>†</sup> |  |
| 585 | Bond lengths, Å | 0.008 |
| 586 | Bond angle, ° | 1.182 |
| 587 | Average B-factor, Å<sup>2</sup> | 86.34 |
| 588 | Mean B factor (Å<sup>2</sup>) | 41.7 |
| 589 | Ramachandran plot** |  |
| 590 | Core, % | 97.5 |
| 591 | Allowed, % | 2.5 |
| 592 | Disallowed, % | 0 |

†Rsym = $(\sum(\text{ABS}(I(h,i) - I(h))) / (\sum(I(h,i)))$ .
‡R-factor = $\sum |F_o - F_c| / \sum |F_o|$ where $F_o$ and $F_c$ are the observed and calculated structure factor amplitudes, respectively.
§R-free is the R-factor calculated with 5% of the reflections chosen at random and omitted from refinement.
<sup>†</sup>rmsd of bond lengths and bond angles from ideal geometry.
\*\*Performed by Procheck.

**Table S3.**
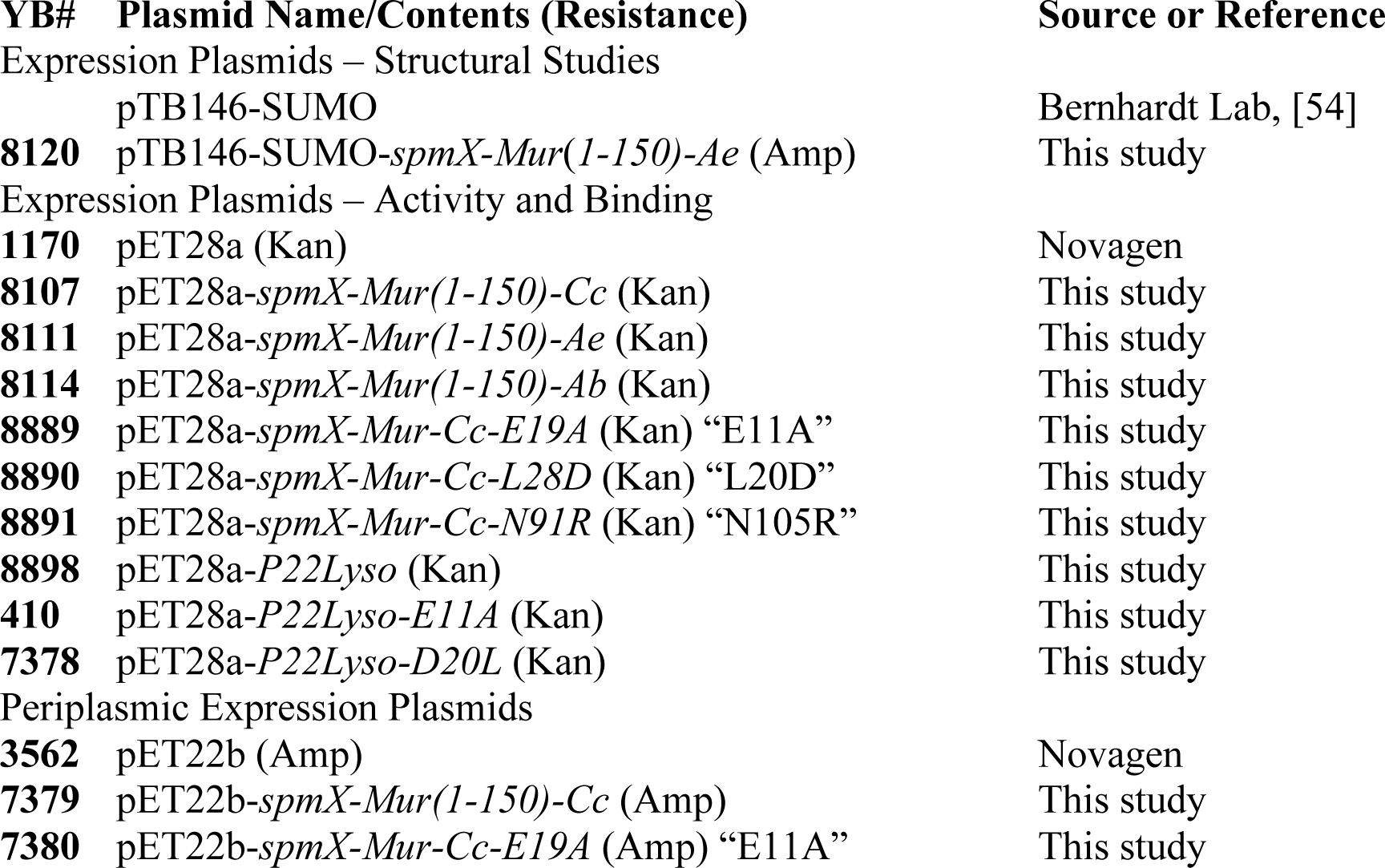
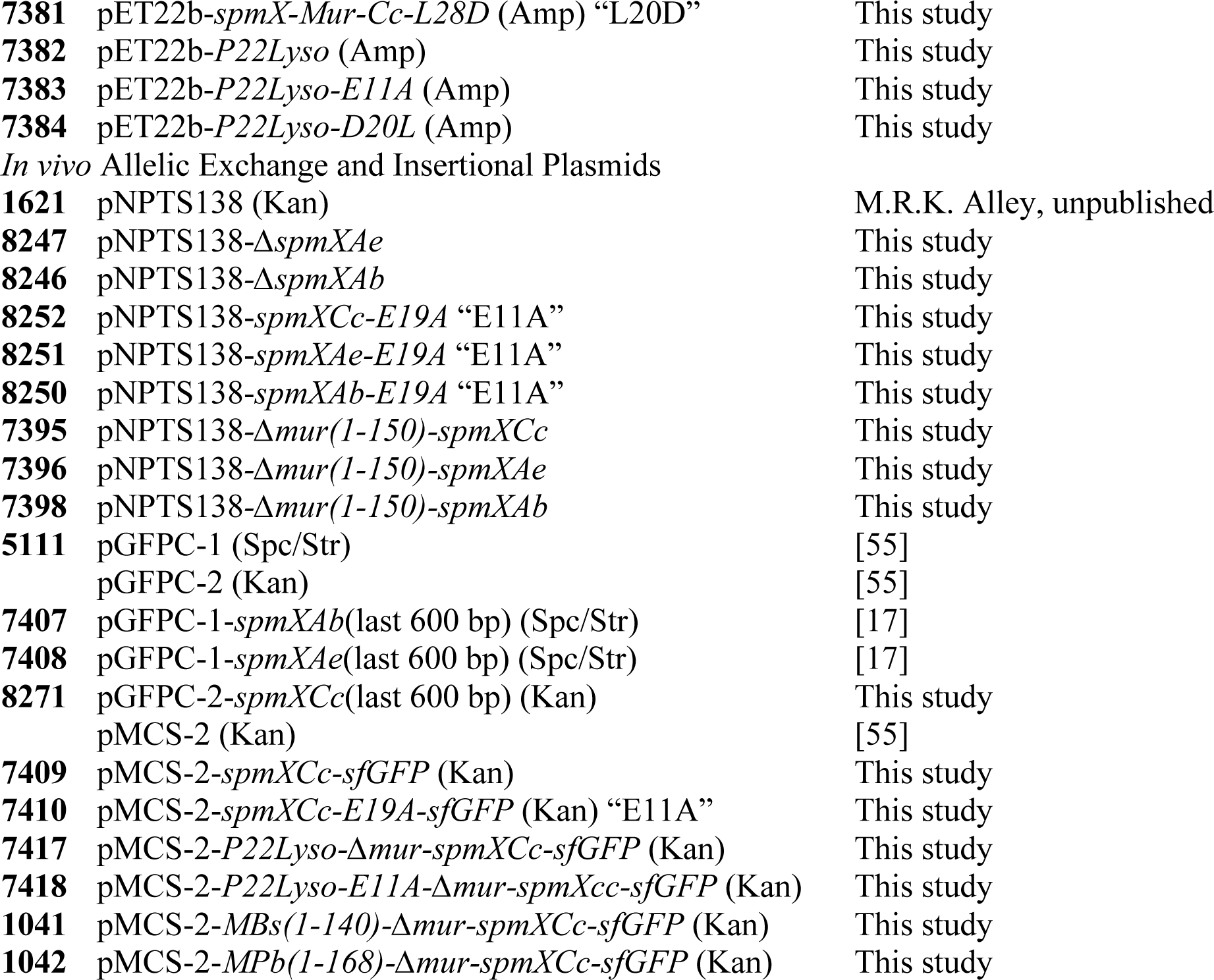
Plasmids

**Table S4.**
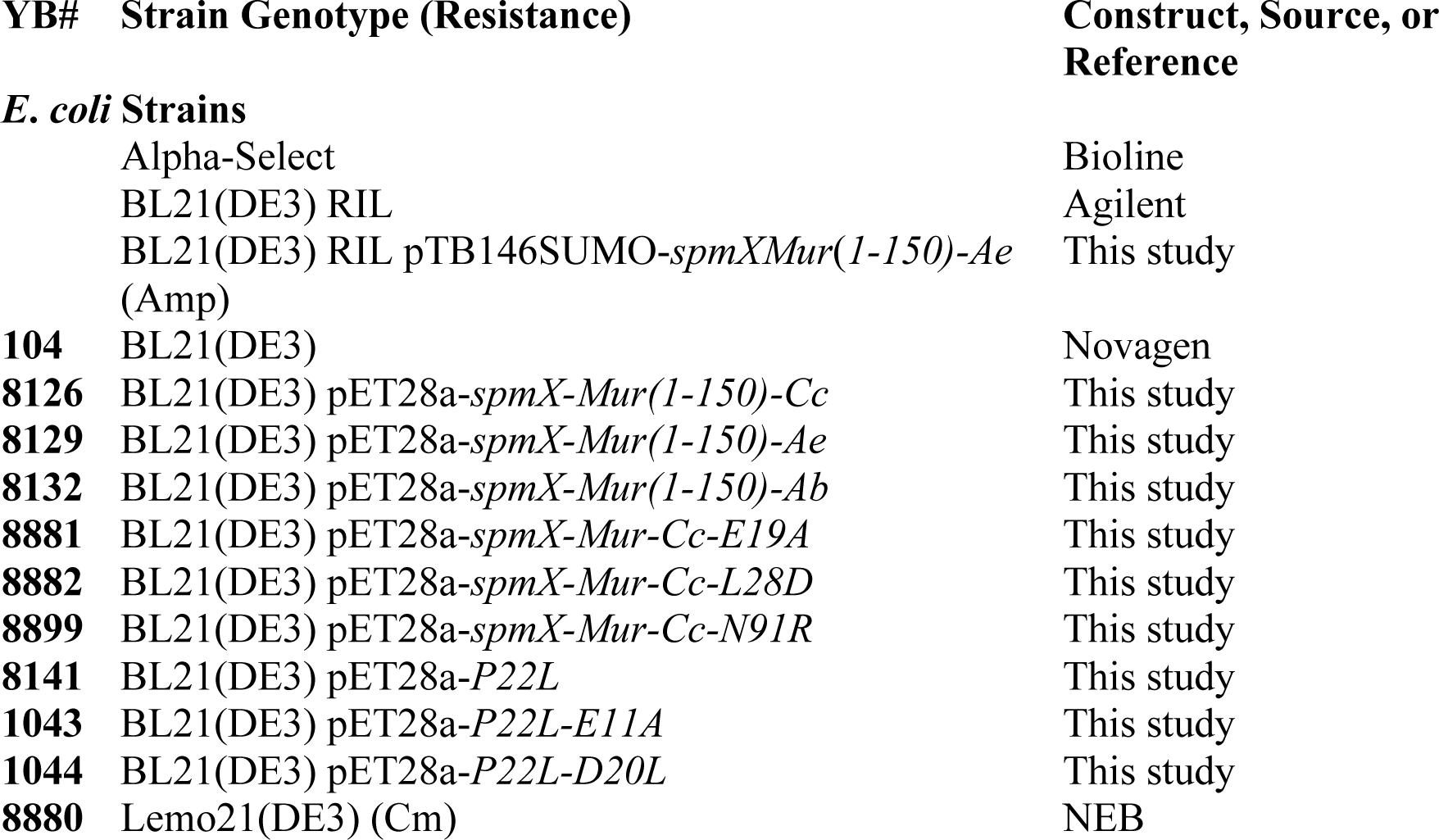
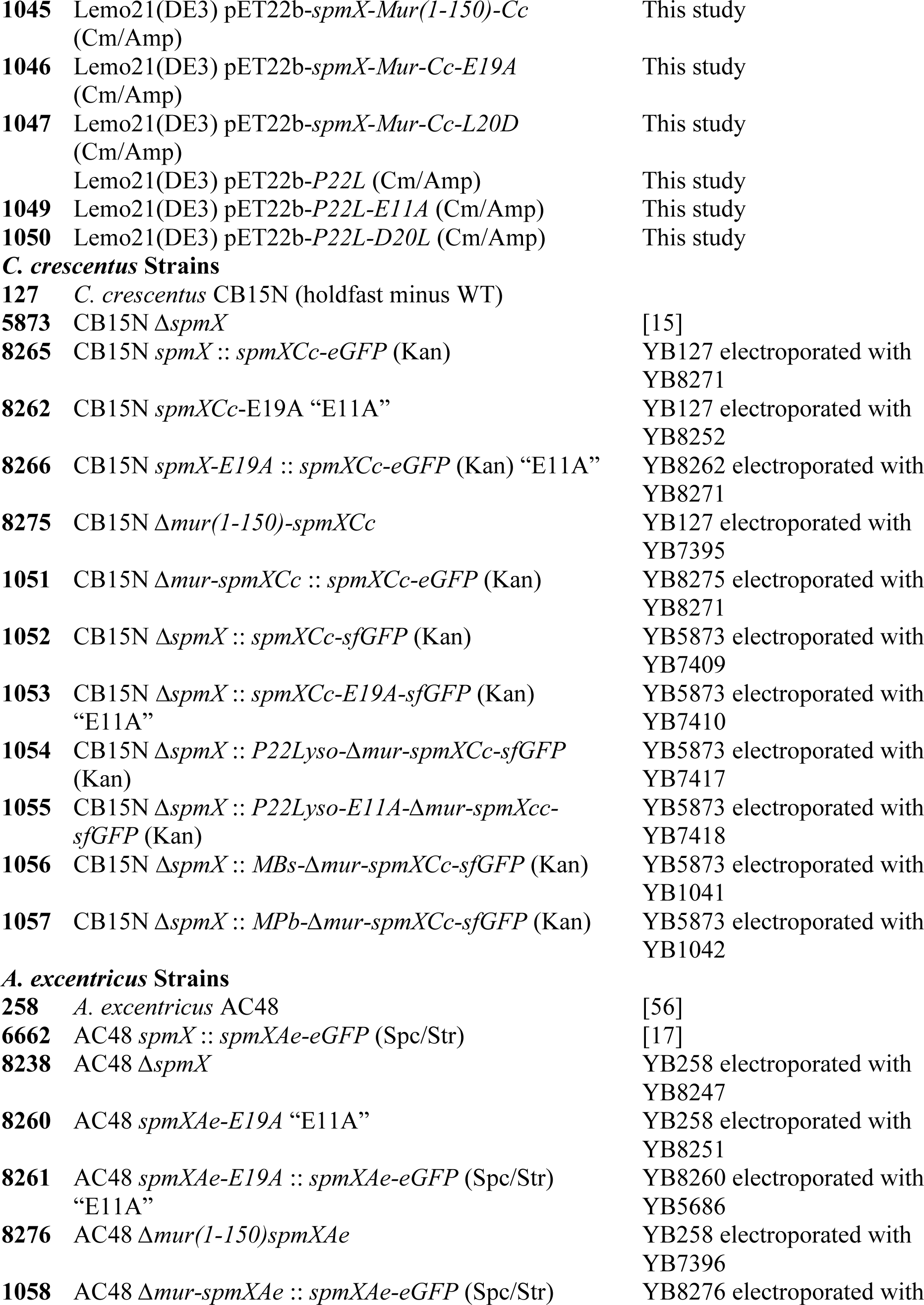
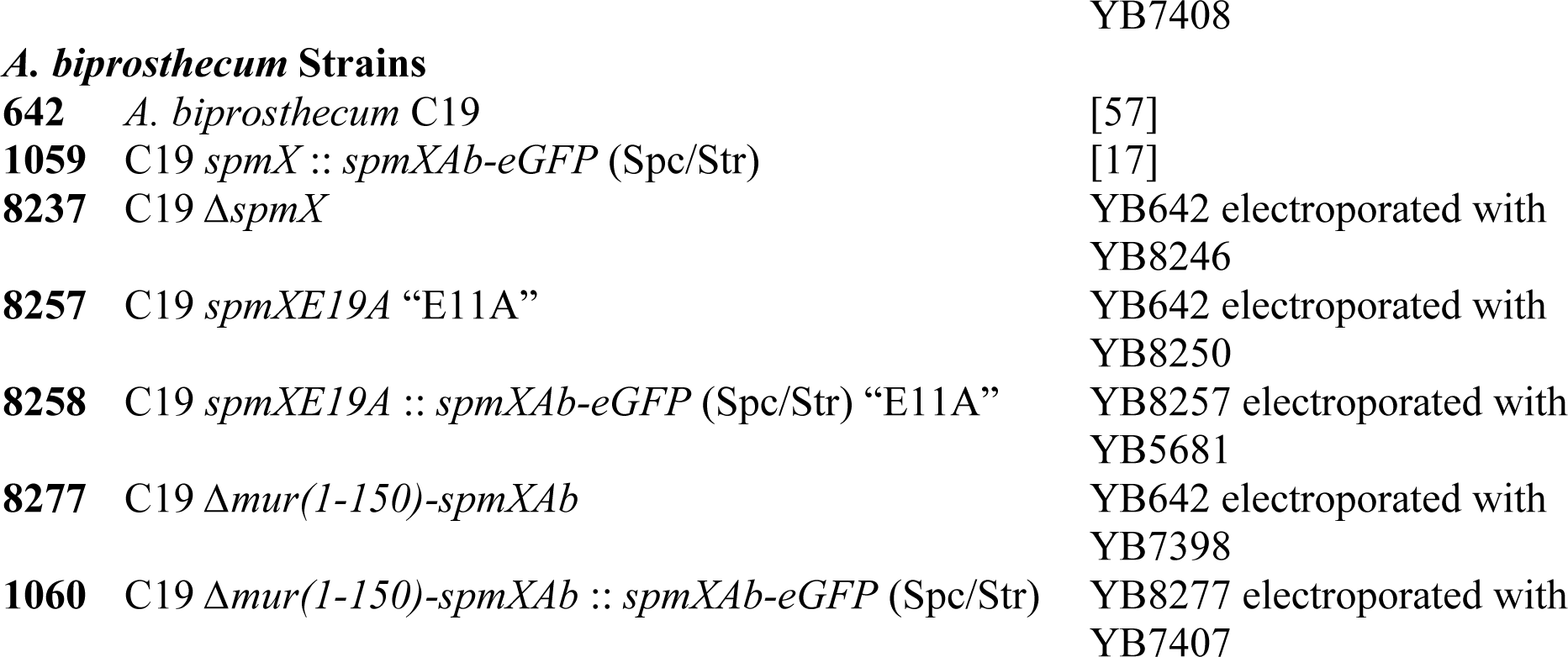
Strains

## Materials and methods

### Bacterial strains and growth conditions

All *C. crescentus*, *A. excentricus*, and *A. biprosthecum* strains used in this study were grown in liquid PYE medium. *C. crescentus* was grown at 30°C and *Asticcacaulis* species at 26°C. Strains were maintained on PYE plates supplemented with antibiotics as necessary (kanamycin 20 μgmL^-1^, gentamycin 5μgmL^-1^, and spectinomycin 100 μgmL^-1^). For microscopy, *C. crescentus* and *A. excentricus* were inoculated from colonies, grown overnight, then diluted back 1:50 and grown for another 3-4 hours before being imaged in mid- to late-exponential phase. *A. biprosthecum* was inoculated from colonies and grown overnight to reach mid- to late-exponential phase for imaging. A detailed list of strains is included as **Table S4**. *E. coli* strains were grown as described in the Methods sections on purification and periplasmic expression.

### Bioinformatics and gene trees

Sequences of the SpmX genes in **Table S1** and members of the GH24 family were retrieved by BLAST searches on the Integrated Microbial Genomes and Microbiomes (IMG/M) database [58] and the National Center for Biotechnology Information (NCBI) “nr” database. Multiple alignments were achieved with MUSCLE [59] and manually adjusted and visualized with Jalview [60]. Sequence conservation of SpmX residues was determined from the multiple sequence alignment of *spmX* alleles using ConSeq [61]. In order to improve visualization of conservation patterns, the ConSeq scores were averaged across a 20-residue sliding window. For estimating bacterial species phylogeny, assembled genome data were obtained from the genome database of the National Center for Biotechnology Information [62]. Amino acid sequences of 37 conserved housekeeping genes were automatically identified, aligned, and concatenated using Phylosift [63]. All phylogenetic reconstruction was performed using MrBayes v3.2.6 [64] to estimate consensus phylogenies and clade posterior probability support values. Sequence substitution was modeled according to a WAG substation model with gamma-distributed rate variation between sites. Trees were visualized and formatted using iTol [65]. The sequence cluster tree was built with NCBI’s Conserved Domain Database tool (CDD). This tool uses reverse position-specific BLAST, a method that compares query sequences to databases of position-specific score matrices and obtains *E-*values, such as in PSI-BLAST [21,22]. WebLogo3 was used to plot the amino acid distribution at each position of the GH motif [53]. To create the alignments for logo generation, 94 T4 lysozyme-like sequences, 20 endolysin/autolysins from the CDD analysis, 60 SpmX muramidase-like sequences (BLAST hits), and 66 SpmX muramidase sequences were simultaneously aligned to T4 lysozyme. Only sequences with unambiguous alignment in the GH motif were included in this analysis.

### Recombinant DNA methods

DNA amplification, Gibson cloning, and restriction digests were performed according to the manufacturer. Restriction enzymes and Gibson cloning mix were from New England Biolabs. Cloning steps were carried out in *E. coli* (alpha-select competent cells, Bioline) and plasmids were purified using Zyppy Plasmid Kits (Zymo Research Corporation). Sequencing was performed by the Indiana Molecular Biology Institute and Eurofins MWG Operon Technologies with double stranded plasmid or PCR templates, which were purified with a DNA Clean & Concentrator kits (Zymo Research Corporation). Chromosomal DNA was purified using the Bactozol Bacterial DNA Isolation Kit (Molecular Research Center). Plasmids were introduced into all *E. coli* strains using chemical transformation according to the manufacturer’s protocols. Plasmids were introduced into *C. crescentus*, *A. excentricus*, and *A. biprosthecum* by electroporation based on previously published studies [66]. Allelic exchange in was achieved with pNPTS138, large genetic insertions with pMCS-2 [55], and eGFP insertional fusions with pGFPC-1 and pGFPC-2 [55].

### Plasmid construction

#### *Expression plasmids* (pTB146SUMO, pET28a, pET22b)

*spmX* gene fragments encoding amino acids 2-150 of SpmX (SpmX-Mur) were amplified from genomic DNA and inserted into linearized expression vectors using Gibson cloning (NEB) according to manufacturers protocols. P22 lysozyme (P22Lyso) was amplified from a synthetic gene strand (Eurofins) for similar construction with Gibson cloning. For pTB147SUMO, the vector was linearized with SapI and XhoI to insert SpmX-Mur-*Ae*. For pET28a, the vector was linearized with NdeI and EcoRI to insert SpmX-Mur-*Cc*, BamHI and XhoI to insert SpmX-Mur-*Ae*, SacI to insert SpmX-Mur-*Ab,* and EcoRI to insert P22Lyso. In all pET28a plasmids, the constructs were cloned in frame with the N-terminal His-tag and a stop codon to eliminate the C-terminal His-tag. For pET22b, the vector was linearized with EcoRI and the C-terminal His-tag was preserved. Point mutants in expression vectors were obtained by using standard quick-change procedures and primers with 3’ single stranded overhangs for increased efficiency.

#### *Integrating plasmids for allelic exchange* (pNPTS138)

For allelic exchange, the desired mutation was engineered into pNPTS138, bracketed by 1 kb up- and downstream of the corresponding genetic region. Integrants were isolated by antibiotic selection and secondary recombination events were selected by sucrose counter-selection using standard procedures. The resulting clones were confirmed by PCR and sequencing isolated genomic DNA.

For genomic deletions of *spmX* in *Asticcacaulis*, pNPTS138 was linearized with EcoRI and codons on either end of the gene were retained to avoid introducing frame-shifts in the surrounding area. Therefore the final gene deletion in *A. excentricus* lacks residues 5-808 and in *A. biprosthecum* lacks residues 5-815. For SpmX∆mur truncations, resides 2-150 were removed in all three species. In all cases pNPTS138 was linearized with EcoRV, except for SpmX*Ab*-E19A, where pNPTS138 was linearized with EcoRI. Point mutations E19A and N91R were integrated into the ∆*spmX* background for ease of clone isolation and included full-length SpmX flanked by 1 kb genetic context. pNPTS138 containing mutated SpmX were constructed using Gibson cloning with fragments on either side of the intended mutation and overlapping primers containing the mutation amplified from genomic DNA. In all cases pNPTS138 was linearized with EcoRV, except for SpmX*Ab*-E19A, where pNPTS138 was linearized with EcoRI.

#### *Integrating plasmids for insertional eGFP fusions* (pGFPC)

The last 600 bp of *spmX* from *C. crescentus* was amplified from genomic DNA and cloned into pGFPC-2 using Gibson cloning.

#### *Integrating plasmids for replacement at the ∆spmX locus* (pMCS-2)

These constructs were designed to allow insertion of various SpmX mutants fused to C-terminal sfGFP into the Δ*spmX* locus in *C. crescentus*. For SpmX-sfGFP and SpmX-E19A-sfGFP, fragments containing 1 kb of genomic DNA upstream of *spmX* and *spmX* or *spmX-E19A* were amplified from existing pNPTS138 constructs and fused to a fragment containing monomeric sfGFP amplified from pSRKKm-Plac-sf*gfp* [67] using Gibson cloning. In the final construct, SpmX and sfGFP are connected with the linker sequence GSAGSAAGSGEF [68]. Chimeras with P22 lysozyme (P22Lyso) and its catalytic mutant were made by Gibson cloning together fragments containing 1kb of upstream genomic DNA, P22Lyso (with no stop codon), SpmX∆mur (residues 151-431) and sfGFP with the same linker. P22Lyso and P22Lyso-E11A were amplified from pET28a plasmids containing these genes. Chimeras with SpmX muramidase from *Brevundimonas subvibrioides* (residues 1-140) and *Parvularcula bermudensis* (residues 1-168) were similarly made with the muramidase fragments amplified from genomic DNA and synthetic gene strands (Eurofins), respectively.

### Production of SpmX-Mur-*Ae* for crystallographic studies

The muramidase domain of SpmX from *A. excentricus* (SpmX-Mur-*Ae*, residues 2-150) was fused to a hexahistidine tag followed by the SUMO cleavage site of the Ulp1 protease (His-SUMO tag) (**Table S3**) [69] and overexpressed in *E. coli* BL21 (DE3) RIL cells. Cells were grown at 37 °C in 2 l of Terrific Broth (BD Biosciences) supplemented with ampicillin (100 μg/ml) until the OD_600nm_ reached 0.8. Production of the recombinant protein was induced by the addition of isopropyl β-D-1-thiogalactopyranoside (IPTG) to 0.5 mM after the culture was cooled to 25°C. Cell growth was continued overnight at 25°C, and cells were harvested by centrifugation. Cell pellets were resuspended in 1/20^th^ volume of buffer A (50 mM Tris-HCl (pH 8.0), 500 mM NaCl, 25 mM imidazole, 10% (vol/vol) glycerol) containing the Complete^TM^ cocktail of protease inhibitors (Roche). Cells were lysed by six passages through a cell disruptor (Constant Systems Limited) at 20 kPsi, and cell debris were pelleted by centrifugation at 40,000 × g for 30 min at 4 °C. The centrifugation supernatant was loaded on a Ni-NTA agarose resin (Qiagen) equilibrated with buffer A. After extensive washing with buffer A, His-SUMO-SpmX-Mur-*Ae* was eluted with a linear 0-100% gradient of buffer B (50 mM Tris-HCl (pH 8.0), 300 mM NaCl, 500 mM imidazole, 10% (vol/vol) glycerol) over 10 column volumes. Peak fractions were pooled, mixed with a 1:100 dilution of a His-tagged Ulp1 (SUMO) protease preparation [70] and dialyzed overnight at 4°C in buffer C (50 mM Tris-HCl (pH 8.0), 300 mM NaCl, 10% (vol/vol) glycerol). Cleavage reactions were passed through Ni-NTA resin to remove free His-SUMO tag and His-Ulp1, and untagged protein was collected in the flow through. Flow-through fractions were concentrated with Amicon Ultra Centrifugal filter units with a molecular weight cutoff of 10 kDa (Millipore) and were injected onto an ENrich^TM^ SEC650 10×300 gel-filtration column (Biorad). SpmX-Mur-*Ae* was eluted with buffer D (25 mM Tris-HCl (pH 8.0), 150 mM NaCl) and again concentrated with Amicon Ultra Centrifugal filter units. Protein concentration was measured using absorbance at 280 nm.

### Protein crystallization and structure determination

High-throughput crystallization trials were performed with a Cartesian PixSys 4200 crystallization robot (Genomic Solutions, U.K.). Hanging drops containing 100 nl of protein (25 or 12.5 mg/ml) and 100 nl of reservoir solution were set up in 96-well Crystal Quick plates (Greiner) and incubated at 20°C. Initial crystal hits were refined manually by setting up hanging drops containing 1 μl of protein (25 or 12.5 mg/ml) and 1 μl of reservoir solution in 24-well plates (Molecular Dimensions) incubated at 20°C. Large needle-shaped crystals (dimensions of about 40 x 40 x 400 μm) were finally obtained for SpmX-Mur-*Ae* in 0.1 M Tris-HCl pH 8.5, 12% PEG 3350, 0.2 M MgCl_2_, at 20°C within 24–48 h. SpmX-Mur-*Ae* crystals were cryoprotected by transfer into 0.1 M Tris-HCl pH 8.5, 13% PEG 3350, 0.2 M MgCl_2_, 10% glycerol, and then flash-frozen in liquid nitrogen. X-ray diffraction data were collected at the European Synchrotron Radiation Facility (ESRF, Grenoble, France) on the ID30a1 (MASSIF-1) beamline [71,72].

Diffraction data were indexed and scaled using the XDS program suite [73]. SpmX-Mur-*Ae* crystals belong to the trigonal space group P3_2_21, with unit cell dimensions of 100.44 x 100.44 x 96.62 Å and three molecules per asymmetric unit. Phase determination was carried out by the molecular replacement method with PHASER [74], using as a search model the structure of the phage P22 lysozyme (PDB entry 2ANX). The molecular replacement solution model was rebuilt de novo using PHENIX [75] to prevent bias from the model.

The structure of SpmX-Mur-*Ae* was completed by cycles of manual building with COOT [76] and addition of water molecules with ARP/wARP [77]. Several cycles of manual building and refinement with REFMAC [78], as implemented in the CCP4 program suite, were performed until R_*work*_ and R_*free*_ converged [79]. Stereochemical verification was performed with PROCHECK [80]. The secondary structure assignment was verified with DSSP [81], with all residues within most favorable or allowed regions of the Ramachandran plot.. Figures were generated with PyMol (http://www.pymol.org). Coordinates of the final refined model were deposited at the Protein Data Bank (PDB, http://www.rcsb.org) and were assigned PDB entry code 6H9D. The data collection and refinement statistics are summarized in **Table S2**).

### Protein production for *in vitro* assays

Fresh BL21(DE3) competent cells (Novagen) were transformed with pET28a constructs containing various muramidase genes with N-terminal His-tags (**Table S3**) and grown overnight in LB with 1% glucose and 50 μg/mL kanamycin. Overnight cultures were diluted 100-fold in LB medium with 1% glucose and 50 μg/mL kanamycin. Typically 500 mL cultures of cells were grown for 1.5-2 hours to an OD600 of 0.6-0.7 and shifted to 20°C. When the OD600 reached 0.8−0.9, the cells were induced with 0.5 mM IPTG. After growing for 4 h at 20 °C, cells were harvested and resuspended in 30 mL lysis buffer (25 mM HEPES pH 7.5, 100 mM NaCl, 20 mM imiadazole, 5 mM BME) with a EDTA-free Protease Inhibitor Mini Tablet (Pierce) and phenylmethanesulfonyl fluoride (PMSF, 1 mM). The 30 mL cell mixture was lysed on ice using a sonicating horn and spun down at 10,000g for 20-30 min. The clarified lysate was loaded onto a 5 ml HiTrap Chelating HP cartridge (GE Healthcare) charged with Ni^2+^ and pre-equilibrated with lysis buffer. After loading, the column was washed with lysis buffer followed by an elution via a 0-100% linear gradient of buffer B (25 mM HEPES pH 7.5, 100 mM NaCl, 500 mM imidazole, 2 mM BME). Muramidase-containing fractions were pooled based on SDS-PAGE analysis and concentrated to 2.5 mL. Imidizole was removed by passing the concentrated fraction over a PD10 desalting column (GE Healthcare) equilibrated with 25 mM HEPES pH 7.5, 100 mM NaCl, 2 mM BME.

### Sacculi preparation and RBB labeling

Sacculi were prepared from all species in the same manner. For a typical 2L prep, cells were grown to an OD of 0.5-0.7 in their respective medium (see bacterial strains and growth conditions) and harvested by centrifugation at 6,000*g* for 20 minutes. *C. crescentus* cells usually required multiple centrifugation steps to collect all the cells. Cells were resuspended in 25 mL water (or PBS for *E. coli*) and added drop-wise into 50 mL of boiling 7.5% SDS under stirring. The mixture was boiled for 30 minutes and then allowed to cool to room temperature. Sacculi were then pelleted by ultracentrifugation at 100,000*g* for 30 minutes at room temperature. The resulting pellets were resuspended in 100 mL pure water, and washed repeatedly until SDS was no longer detected in the supernatant. The pellet was confirmed to be clear of SDS by mixing 0.2 mL of the supernatant with 1 uL 0.5% methylene blue, 0.1 mL 0.7M NaPO_4_ pH 7.2, and 0.6 mL chloroform and checking to make sure that, after vortexing and allowing to settle, the solution had an upper blue phase and a lower clear phase [82]. At this point, the pellets were resuspended in 10 mL PBS with 20 mM MgSO_4_, 250 U/uL Pierce Universal Nuclease (Thermo Fisher Scientific), and 10 mg/mL amylase (Sigma). The mixture was incubated at 37°C for 1-4 hours. Afterwards, 10 mg/mL trypsin and 10 mM CaCl_2_ was added and the mixture incubated overnight at 37°C. 800 uL of 15% SDS were then added to the mixture and it was brought to a boil for about 10 minutes and allowed to cool to room temperature. The sample was then pelleted (100,000*g*, 30 min, room temperature) and resuspended in 4-5 wash steps until SDS was no longer detected. The final pellet was then resuspended in 2 mL water and added to 0.8 mL 0.2M remazol brilliant blue (Sigma), 0.4 mL 5M NaOH, and additional water to 8 mL. The mixture was incubated, shaking, overnight at 37°C. After neutralizing the solution with 0.4 mL 5M HCl. The mixture was then pelleted (21,000*g,* 20 minutes, room temperature), and resuspended in water until the supernatant became clear.

To calibrate the concentration of RBB-labeled sacculi for dye-release assays and peptidoglycan-binding assays, activity curves with Hen Egg White Lysozyme (Sigma) were produced using different dilutions of the RBB-labeled sacculi. The RBB-labeled sacculi were used at the dilution that resulted in an A595 of 0.5 when 5 μL of the RBB-labeled sacculi were incubated with 4 uM HEWL.

### Peptioglycan-binding assays

5μL of calibrated RBB-labeled sacculi were incubated with 1 μM of purified protein (protein constructs used are shown in Fig. S4) in PBS pH 7.4 to a final volume of 50 μL for 30 minutes at 37°C and then pelleted (16,000*g*, 20 minutes). Fractions were separated and the pellet resuspended in 50 μL PBS. 10 μL of each fraction was loaded onto Any kD Mini-PROTEAN TGX Precast Protein Gels (BioRad) to visualize whether the protein associated with the insoluble sacculi fraction. BSA (Sigma) was used at 1 μM as a negative control.

### Remazol brilliant blue dye-release assays

Methods were adapted from [32,83]. Assays were carried out in 25-μL reactions using 25 mM HEPES pH 7.5, 100 mM NaCl and 5 μL of calibrated RBB-labeled sacculi. Enzymes were added at various concentrations (see Figs. 5 and S5) and incubated overnight at 37°C. Reactions were then centrifuged for 20 minutes at 16,000*g,* and the supernatant carefully separated from the pellet. Final values in Fig. 5 and S5 are normalized against absorbances measured for reactions with HEWL that were run in tandem for every measurement to correct for differences in different sacculi preparations.

### Fluorescence microscopy and image analysis

Fluorescence imaging was done using an inverted Nikon Ti-E microscope using a Plan Apo 60X 1.40 NA oil Ph3 DM objective with a GFP/Cy3 filter cube and an Andor DU885 EM CCD camera. Images were captured using NIS Elements (Nikon). Cells were mounted on 1% (w/v) agarose pads made with PYE (or PBS, in the case of *E. coli*) for imaging. In general, the fluorescent channel of each image was background subtracted and a Gausian Blur filter was applied using Fiji [84]. Quantification of stalk morphotypes and stalks with multiple foci was done by hand using Fiji tools. Quantification of fluorescence data was achieved using MicrobeJ [85]. Mean stalk intensity was measured in *A. biprosthecum* cells by using Fiji to draw line ROIs that did not overlap with the focus at the base of the stalk, measuring mean fluorescence along the ROI. Figures and statistics were performed using GraphPad Prism version 8.00 for Mac, GraphPad Software, La Jolla California USA, www.graphpad.com.

### Western blots

Strains were grown to saturation (overnight for *C. crescentus* and *A. excentricus*, usually 48 hours for *A. biprosthecum*). OD600 was determined and cells were collected at a normalized density of OD600 = 1/1mL. 1 mL of each normalized culture was pelleted, resuspended in 100 uL water, and prepared for analysis using standard procedures using SDS-PAGE, transfer, and western blotting. 10 μL of each sample was loaded onto Any kD Mini-PROTEAN TGX Precast Protein Gels (BioRad). The JL-8 monoclonal GFP antibody (Clontech) was used as the primary antibody and Goat Anti-mouse HRP (Pierce) was used for the secondary antibody. Transferred blots were visualized with SuperSignal West Dura Extended Duration HRP substrate (ThermoFisher Scientific) using a Bio-Rad Chemidoc.

### Periplasmic expression in *E. coli*

Fresh Lemo(DE3) competent cells (NEB) were transformed with pET22b constructs containing various muramidase genes with N-terminal H-tags (**Table S3**) and plated. Lemo21(DE3) carries a rhamnose-inducible copy of LysY that inhibits T7 polymerase and allows for tunable dampening of expression of toxic products. We could not transform expression strains BL21(DE3) or Tuner(DE3) with the pET22b-P22Lyso construct, but were able to isolate a few transformants carrying this construct using Lemo21(DE3) cells under high rhamnose repression (2 mM). P22Lyso-D20L, and all the SpmX-Mur-*Cc* constructs, efficiently transformed into all expression strains tested, and could be carried by Lemo21(DE3) without rhamnose.

In the case of pET22b-P22Lyso, where cells lyse from leak, cell cultures were grown directly from colonies in the presence of 5 mM rhamnose and monitored over time. Figure 4A shows the same treatment for all tested constructs. For testing induction of CCM and P22Lyso-D20L, colonies were grown overnight in LB with 100 ug/mL carbenicillin and 30 μg/mL chloramphenicol. Overnight cultures were diluted 50-fold in LB medium with 100 ug/mL carbenicillin and 30 μg/mL chloramphenicol. In experiments cases rhamnose was added at specified concentrations. Typically 4 mL cultures of cells were grown for 1-1.5 hours to an OD600 of 0.3-0.4, induced with 400 uM IPTG, and shifted to 20°C.

#### Growth curves and live-dead staining

Optical densities were measured over time and cells were routinely checked for lysing by microscopy using standard procedures. Briefly, 1 uL of a 1:1 mixture of solutions A and B from a LIVE/DEAD *Bac*Light Bacterial Viability Kit (ThermoFisher Scientific) was directly added to 100 uL of cells diluted 1:10 in PBS. Cells were visualized on 1% agar pads made with PBS using the methods described in microscopy.

#### Periplasmic expression levels

After growing for 4 h at 20 °C, OD600 was determined and cells were collected at a normalized density of OD600 = 1/1mL. One mL of the normalized sample was pelleted at 4000g for 15 min and the pellet resuspended in 250 uL 20% sucrose, 1 mM EDTA, 30 mM TRIS pH 8 at room temperature. The sample was mixed gently by rotation at room temperature for 10 minutes before being spun down at 13,000g for 10 minutes. The supernatant was carefully removed and the pellet rapidly suspended in 250 uL ice cold pure water. The sample was mixed gently by rotation at 4°C for 10 minutes before being spun down at 13,000g at 4°C. The supernatant (periplasmic fraction) and pellet (cell fraction) were then separated and prepared for analysis using standard procedures using SDS-PAGE, transfer, and western blotting. Blots were incubated with His-Probe Antibody (H-3) sc-8136 HRP (Santa Cruz Biotechnology) and visualized with SuperSignal West Dura Extended Duration HRP substrate (ThermoFisher Scientific) using a Bio-Rad Chemidoc.

## Supplemental Figure Legends

**Figure S1. Alignments of SpmX muramidases. Related to Figure 1.** Aligned protein sequences of the muramidase domains from genes listed in **Table S1**, a subset of which were used for the gene tree in **Fig. 1.** Key positions discussed in text are marked with a black dot and annotated with T4L identities.

**Figure S2. Relationships of enzyme families within the lysozyme superfamily. Related to Figures 1 and 2.** Sequence cluster tree diagram made using NCBI’s Conserved Domain Database tool (CDD) demonstrating the inferred relationship of various lysozyme families with structural homology but no sequence similarity. This tool uses reverse position-specific BLAST, a method that compares query sequences to databases of position-specific score matrices and obtains *E-*values, such as in PSI-BLAST [21,22]. SpmX from *C. crescentus* is marked in red type; bacteriophages discussed in the text are emphasized in bold, black type. The tree was formatted using iTol [65].

**Figure S3. Binding assays of SpmX to peptidoglycan from various species. Related to Figure 4.** Coomassie brilliant blue stained SDS–PAGE gel of supernatant (S) and pellet (P) fractions from peptidoglycan binding assays. In brief, SpmX protein constructs from three different species and of two different lengths were incubated in PBS with or without sacculi (peptidoglycan) isolated from the three different species. Peptidoglycan is insoluble and can be pelleted out of solution. As shown here, peptidoglycan from all three species pulled down the SpmX protein constructs.

**Figure S4. *In vitro* activity and periplasmic expression/activity of SpmX and various mutants. Related to Figure 4.** Active enzymes release peptidoglycan monomers covalently-bound to RBB into the supernatant that are detected by absorbance at 595 nm. (**A**) RBB assays with SpmX-Mur-*Cc-*E11A, the catalytic mutant. Due to poor expression and instability, SpmX-Mur-E11A was not concentrated to higher than 12 μM and its activity is compared to WT SpmX at 12 μm. Error bars are ± standard deviation from two replicates. (**B**) RBB assays with SpmX-Mur-*Cc-*L20D, the mutant that restores the ancestral D20. **(C)** Final ODs of cultures expressing various P22Lyso and SpmX-Mur-*Cc* constructs after four hours of induction. P22Lyso-D20L showed signs of spheroplast formation and cell lysis but the other tested constructs did not. Compare to growth curves and microscopy shown in Fig. 4CD. **(D)** SpmX-Mur-*Cc* is expressed at similar levels in the periplasm as P22Lyso-D20L. Attempts at expressing SpmX-Mur-*Cc* more slowly in M9G medium at lower temperatures increased periplasmic levels but did not result in lysis. (**E**) RBB assays on *E. coli* sacculi isolated from Lemo21(DE3) using purified P22 lysozyme, P22 lysozyme D20L mutant, *C. crescentus* SpmX muramidase, and SpmX muramidase L20D mutant. Lines are drawn to help guide the eye toward basic trends. Normalization is to Hen Egg White Lysozyme (HEWL) assays (see methods).

**Figure S5. Quantification of fluorescence and morphology data of SpmX mutant strains. Related to Figure 5. (A)** Histograms of mean total cell fluorescence and integrated focal intensity of *C. crescentus* cells expressing fluorescently tagged WT (n = 1057 cells) and E11A (n = 792 cells) SpmX. Insets display the mean of these measurements with the standard deviation as error bars. Asterisks indicate p < 0.001 by Mann-Whitney U test. **(B)** Histograms (as in **A**) of mean total cell fluorescence and integrated focal intensity of *A. biprosthecum* cell expressing fluorescently tagged WT (n = 494 cells) and E11A (n = 462 cells) SpmX. Far right panel: boxplot of median and quartiles with whiskers indicating minimum and maximum values of mean fluorescence intensity of stalks in *A. biprosthecum* expressing WT or E11A (n = 10 stalks each strain). **(C)** Histogram (as in **A**) of mean total cell fluorescence and integrated focal intensity of *A. excentricus* expressing fluorescently tagged WT (n = 750 cells) and E11A (n = 693 cells) SpmX. Far right panel: quantification of the frequency of cells with multiple SpmX foci in the stalks of *A. excentricus* WT and E11A mutant. Chi-squared p < 0.001, n = 600 cells. **(D)** Quantification of the stalked morphotypes from *A. biprosthecum* WT and *spmX-E11A* mutant from cells without eGFP tags. Chi-squared p < 0.001, n = 500 cells. **(E)** Panels show bands from two independent blots using samples normalized by OD. Lanes have been cropped and rearranged to aid interpretation. In all cases, the primary antibody is directed against the C-terminal GFP fusion.

